# Defining the *Mycobacterium tuberculosis* Pangenome and Suggestions for a New Composite Reference Sequence

**DOI:** 10.1101/2025.11.03.686283

**Authors:** Poonam Chitale, Elissa Ocke, Aubrey R. Odom, Howard Fan, Alexander Henoch, Karla Vasco, Emily C. Fogarty, Courtney Grady, Alexander D. Lemenze, Pradeep Kumar, Shannon Manning, A. Murat Eren, W. Evan Johnson, David Alland

## Abstract

*Mycobacterium tuberculosis* (Mtb) causes tuberculosis (TB), a global disease with diverse clinical and microbiological manifestations. Studies into the biological causes of this phenotypic diversity have been largely limited to a few reference strains. A pangenome approach is likely to provide new insights. Pangenomic tuberculosis studies have been limited the availability of only fragmented genome sequences and error-prone reference genomes. We used a de novo assembly pipeline that generates extremely complete and accurate whole genome sequences to generate 50 closed Mtb genomes across all seven major lineages. We identified 3,377 core gene clusters and 379 accessory clusters. Analysis showed multi-copy core clusters were largely due to gene fragmentation (76%), paralogs (12%), nearly identical gene duplications (4%), or combinations (8%). Sixteen hypervariable regions (HVRs) were identified, including novel paralogs and variable PE/PPE genes. We consolidated these findings into a Pangenome Gene Reference Resource (PGRR) for precision alignment. Our study demonstrates the closed nature of the Mtb pangenome, with most variation in accessory genes and HVRs. The PGRR provides a foundation for improved drug/vaccine target discovery and highlights the need to move beyond the commonly used H37Rv strain to study Mtb genetic and phenotypic diversity.

**IMPORTANCE:** Tuberculosis (TB), caused by *Mycobacterium tuberculosis*, affects millions globally. Genetic differences among Mtb strains have been difficult to resolve due to incomplete genome references. We sequenced and analyzed complete genomes of 50 Mtb strains from all lineages, identifying 16 hypervariable regions and 3,498 core gene clusters whose diversity mostly stemmed from gene fragmentation, paralog duplication and deletion events and differences in the PE/PPE gene family representation. These differences may explain many of the varied clinical manifestations of TB. We created Pangenome Gene Reference Resource to unify genetic data for precise comparison studies to aid in developing new drugs vaccines and other interventions against this disease.

## INTRODUCTION

Tuberculosis (TB) infects an estimated 10.6 million people and causes 1.6 million deaths annually [1]. TB is caused predominantly by *Mycobacterium tuberculosis sensu stricto* (Mtb), and *Mycobacterium africanum* [2, 3]. Starting in the 1950s, the introduction of TB chemotherapy drove Mtb strains to acquire an expanding repertoire of drug resistance mutations, with several predominant mutations directly related to drug-target interactions and others to changes in metabolic or other cellular functions with more subtle but still measurable effects on drug resistance [4–12]. Slower progress has been made determining the genetic causes of more complex phenotypes such as transmissibility, virulence, or host range. Genetic studies of these phenotypes are likely to benefit from whole genome analysis of large Mtb populations that includes much of the species genetic diversity. The analyses performed so far have relied on error-prone draft genomes and a limited pool of reference strains—most commonly H37Rv, which itself contains annotated errors [13–15] and is further confounded by the highly repetitive PE/PPE family (nearly 10% of the Mtb genome) that resists accurate assembly and annotation [16, 17].

Recent advances in long-read sequencing and high-fidelity assembly permit generation of complete, error-free Mtb genomes [13] which can be leveraged by pangenomic analysis to reveal the entire gene repertoire of the species. The information gained regarding differences in gene representation can potentially explain much of the phenotypic diversity observed in TB including determinants of drug resistance, host-pathogen interactions, virulence and transmissibility. Prior pangenome analyses have been limited by reliance on short-read draft genomes, reference bias, and incomplete representation of all major global lineages [18–21]. Furthermore, critical regions— especially PE/PPE gene loci—are rarely resolved. High-quality, lineage-diverse, closed genomes are therefore essential to map the true diversity of the Mtb pangenome.

Here, we present a comprehensive pangenome based on 50 high-fidelity, completely closed Mtb genomes, representing all seven major lineages. We include an *M. canettii* strain to root evolutionary analyses but exclude it from the pangenome proper. We also introduce the Pangenome Gene Reference Resource (PGRR)—a consolidated reference for all genes and variants found—enabling researchers to precisely align, compare, and annotate new TB genomes.

## METHODS

Details on bacterial strains used, pyazinamidase activity determination, culture, DNA extraction, Nanopore and Illumina whole genome sequencing along with assembly and annotation, lineage SNP and indel identification, phylogenetic and pangenome analysis, and the Pangenome Gene Reference Resource are provided in “Supplementary Methods”.

## RESULTS

### Strain collection and sequencing analysis

We analyzed 50 highly accurate Mtb genomes generated with Bact-Builder, a bioinformatics pipeline which uses a consensus-based approach to generate highly accurate and complete closed bacterial genome sequences [13] that span all seven established lineages to describe the Mtb pangenome (Table 1). For comparison and annotation purposes, our analysis included well characterized reference or laboratory strains H37Rv (TMC102 and NR-123), CDC1551, Erdman and HN878. The remaining strains were obtained from either the Tropical Disease Research (TDR)-TB strain bank established by the World Health Organization Special Program for Research and Training in Tropical Disease [22] or a collection of 20 ‘reference’ strains representing all seven lineages described previously (Table S1) [22, 23]. The average genome size was 4,413,145 ± 37,039 (mean ± SD) base pairs (bp) with an average GC content of 65.61% (Table 1). Annotated genomes had 4,212 to 4,589 coding sequences (CDS) with an average of 4,266.62 ± 50.8, of which 829 to 916 were hypothetical CDS’s with an average of 855.24 ± 14.1 (Table 1).

**Table 1.**
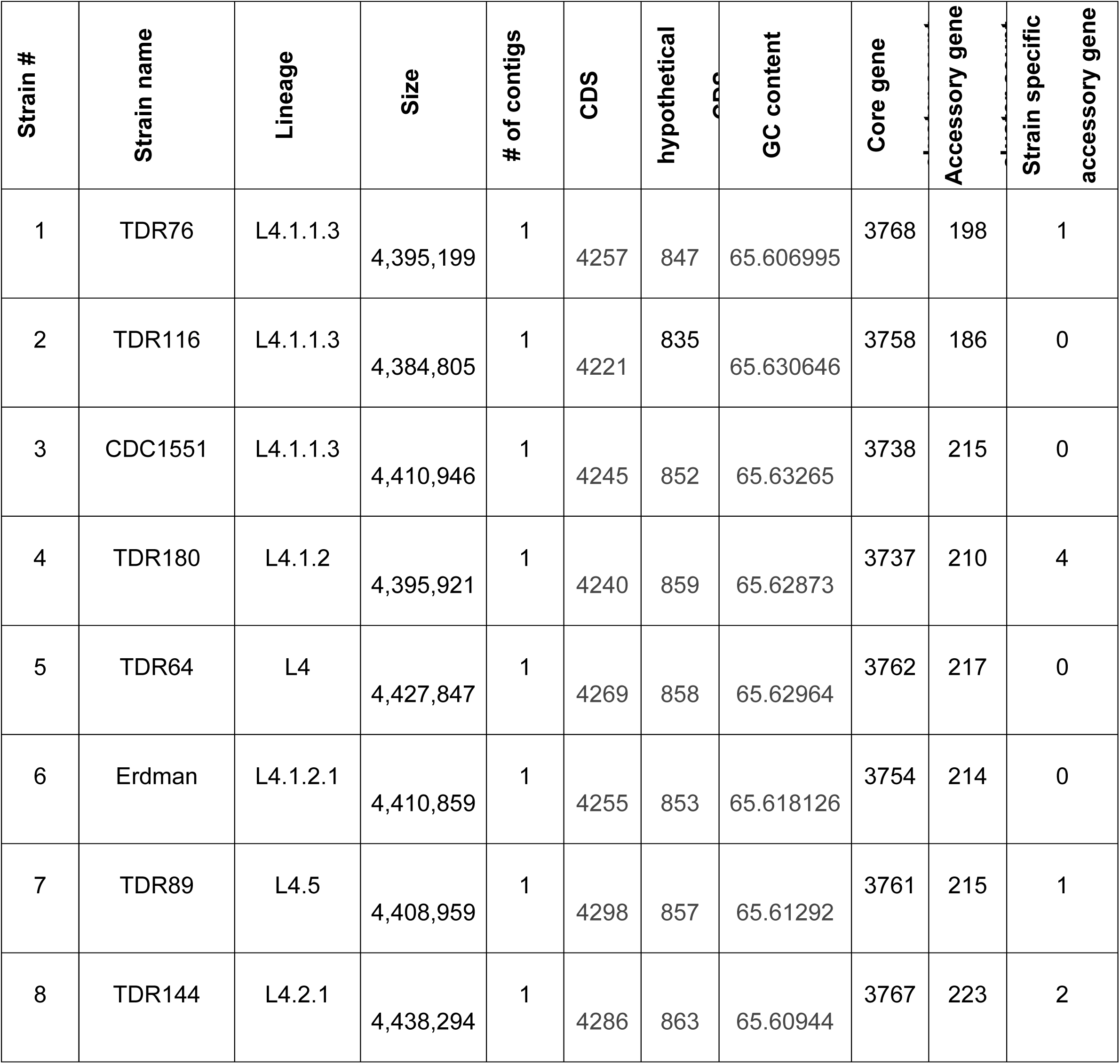
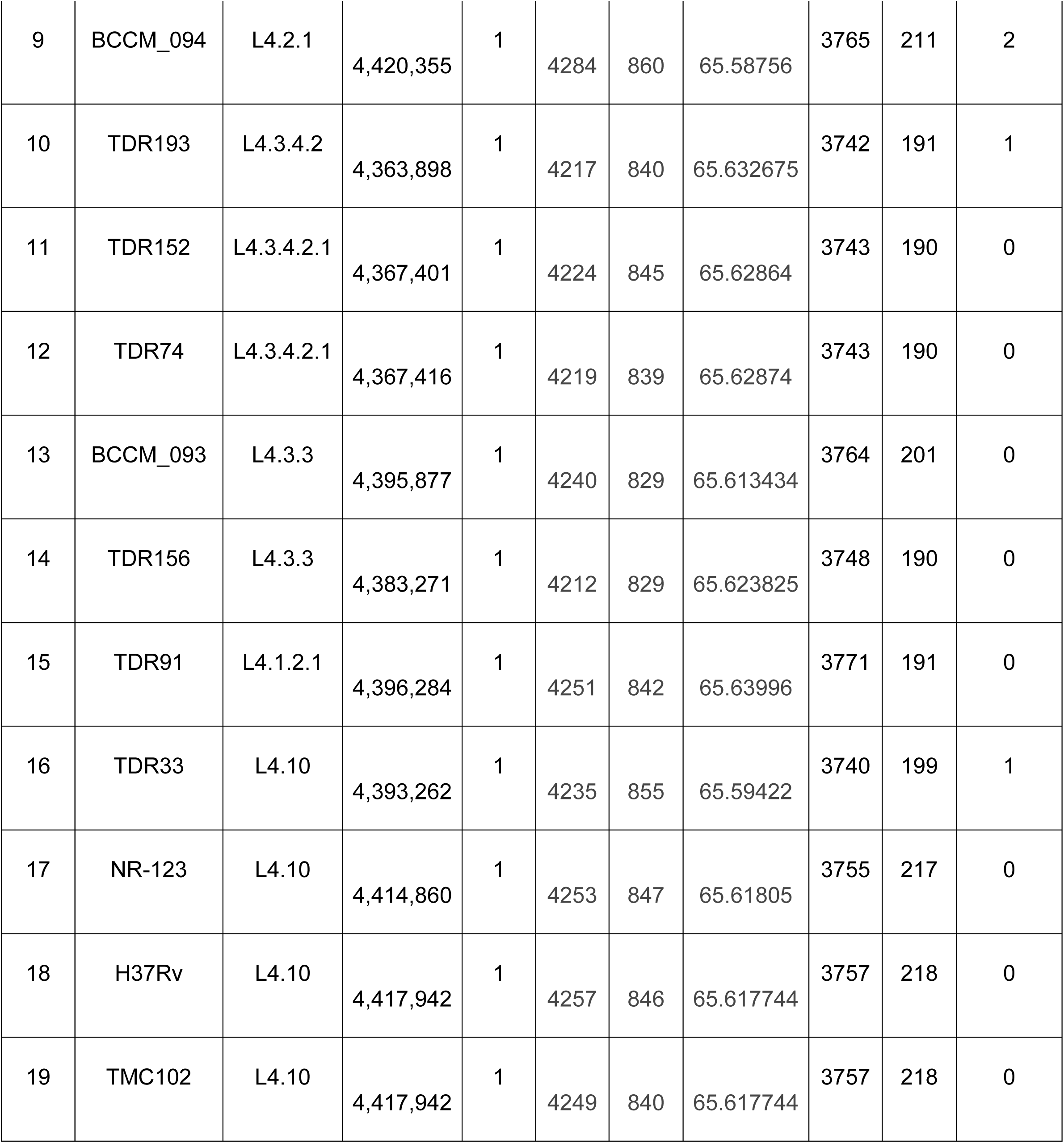
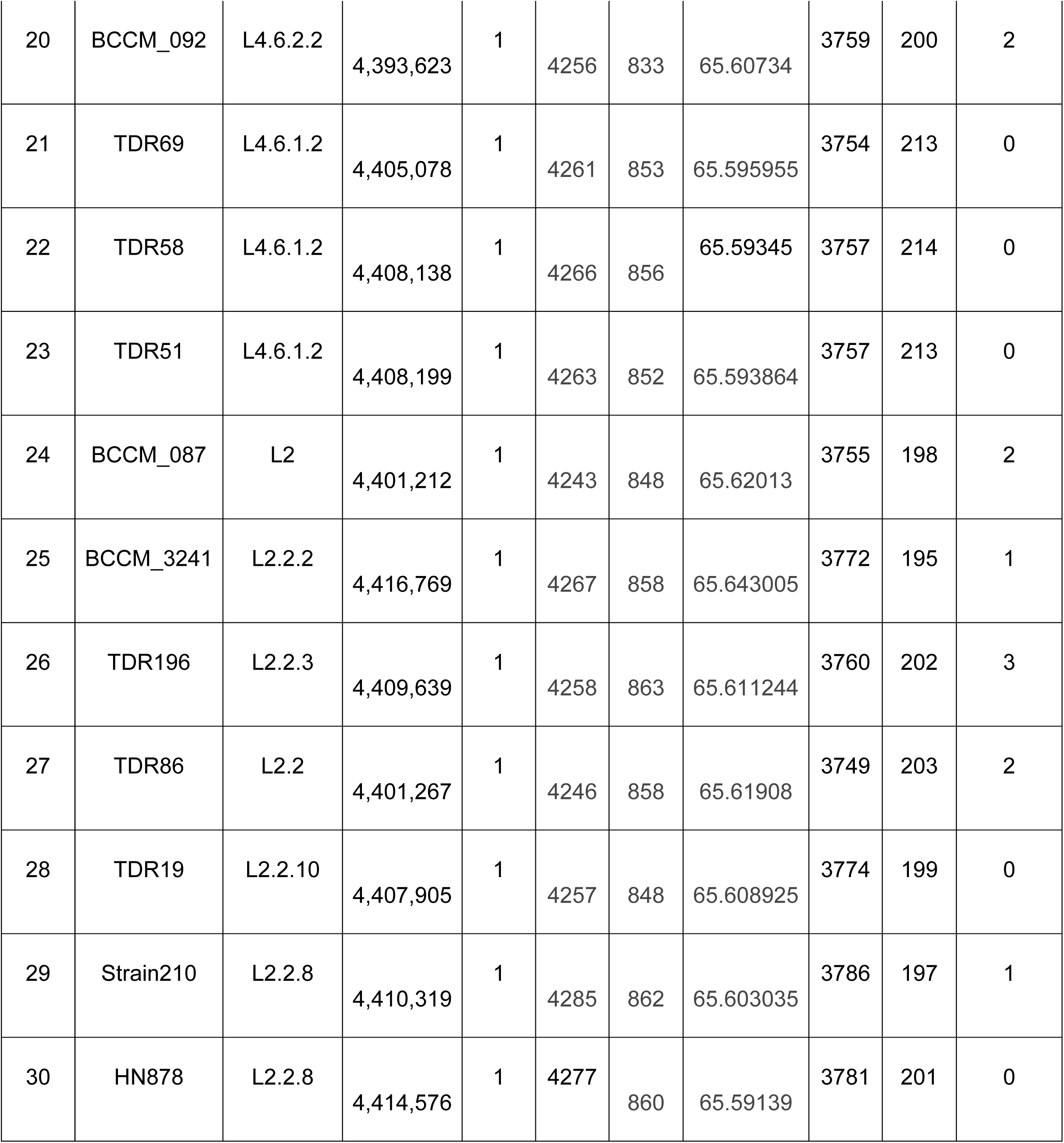
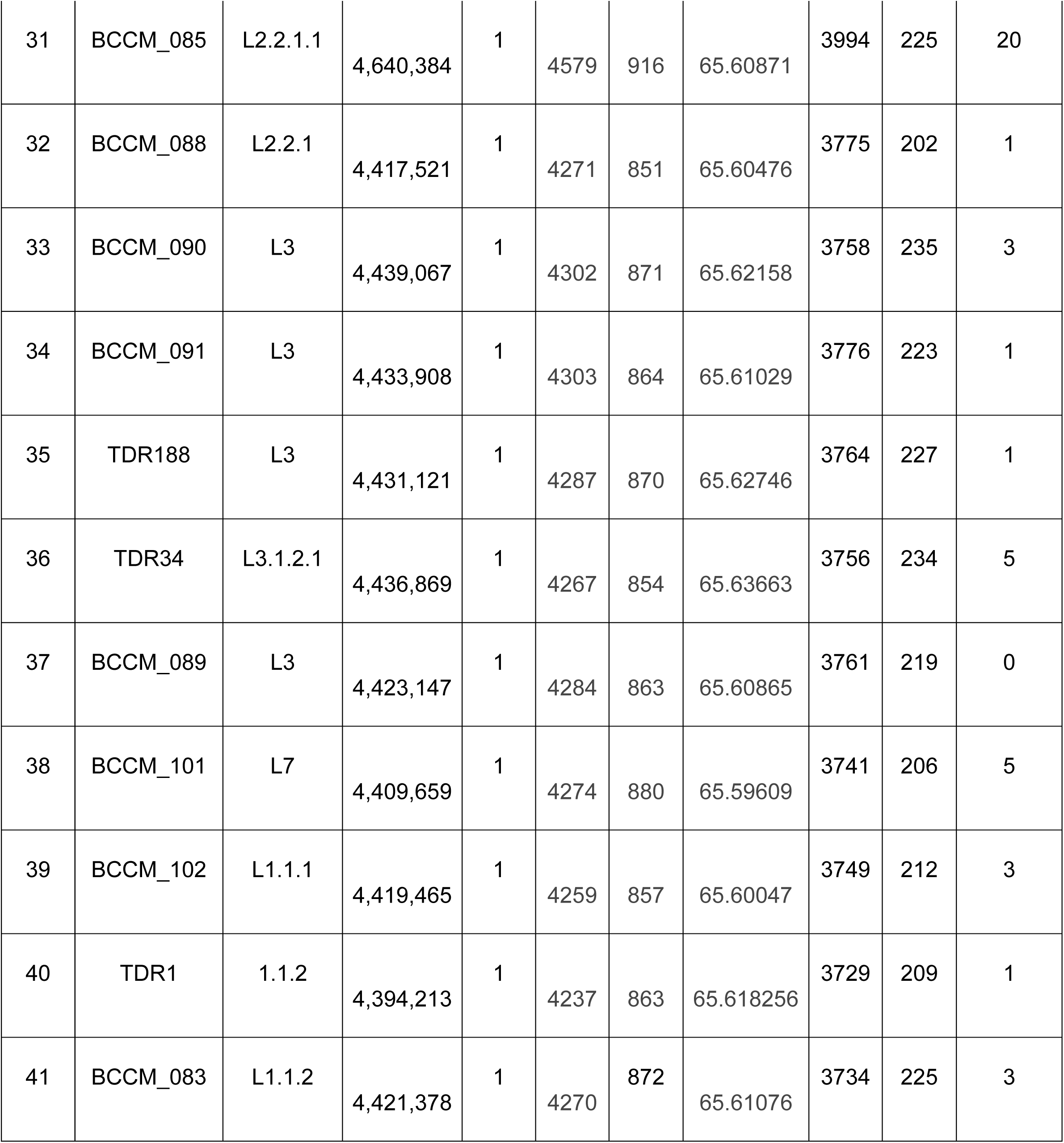
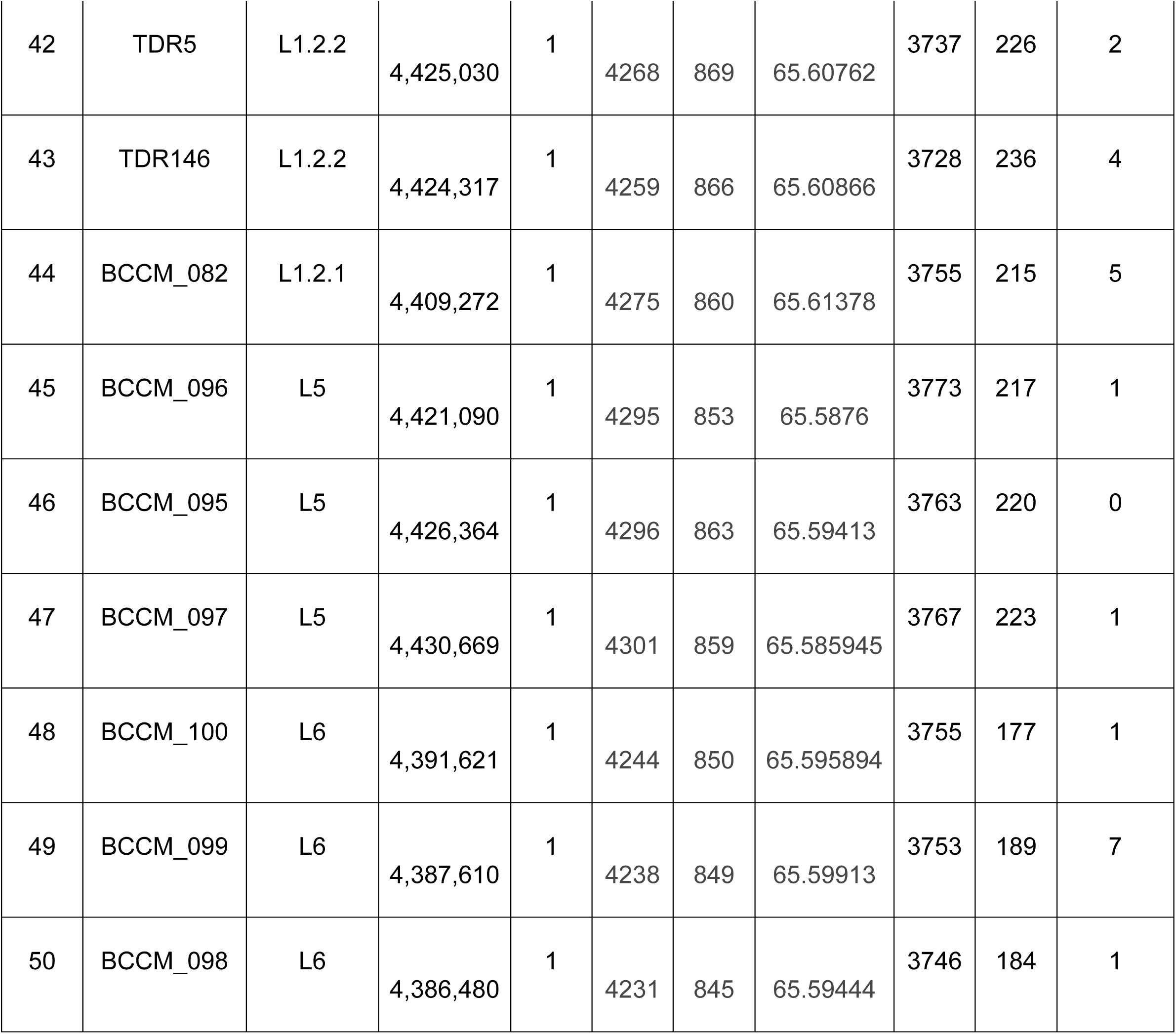
Pangenome strain lineages and genome size.

| **Strain #** | **Strain name** | **Lineage** | **Size** | **# of contigs** | **CDS** | **hypothetical** | **GC content** | **Core gene** | **Accessory gene** | **Strain specific accessory gene** |
| --- | --- | --- | --- | --- | --- | --- | --- | --- | --- | --- |
| 1 | TDR76 | L4.1.1.3 | 4,395,199 | 1 | 4257 | 847 | 65.606995 | 3768 | 198 | 1 |
| 2 | TDR116 | L4.1.1.3 | 4,384,805 | 1 | 4221 | 835 | 65.630646 | 3758 | 186 | 0 |
| 3 | CDC1551 | L4.1.1.3 | 4,410,946 | 1 | 4245 | 852 | 65.63265 | 3738 | 215 | 0 |
| 4 | TDR180 | L4.1.2 | 4,395,921 | 1 | 4240 | 859 | 65.62873 | 3737 | 210 | 4 |
| 5 | TDR64 | L4 | 4,427,847 | 1 | 4269 | 858 | 65.62964 | 3762 | 217 | 0 |
| 6 | Erdman | L4.1.2.1 | 4,410,859 | 1 | 4255 | 853 | 65.618126 | 3754 | 214 | 0 |
| 7 | TDR89 | L4.5 | 4,408,959 | 1 | 4298 | 857 | 65.61292 | 3761 | 215 | 1 |
| 8 | TDR144 | L4.2.1 | 4,438,294 | 1 | 4286 | 863 | 65.60944 | 3767 | 223 | 2 |
| 9 | BCCM_094 | L4.2.1 | 4,420,355 | 1 | 4284 | 860 | 65.58756 | 3765 | 211 | 2 |
| 10 | TDR193 | L4.3.4.2 | 4,363,898 | 1 | 4217 | 840 | 65.632675 | 3742 | 191 | 1 |
| 11 | TDR152 | L4.3.4.2.1 | 4,367,401 | 1 | 4224 | 845 | 65.62864 | 3743 | 190 | 0 |
| 12 | TDR74 | L4.3.4.2.1 | 4,367,416 | 1 | 4219 | 839 | 65.62874 | 3743 | 190 | 0 |
| 13 | BCCM_093 | L4.3.3 | 4,395,877 | 1 | 4240 | 829 | 65.613434 | 3764 | 201 | 0 |
| 14 | TDR156 | L4.3.3 | 4,383,271 | 1 | 4212 | 829 | 65.623825 | 3748 | 190 | 0 |
| 15 | TDR91 | L4.1.2.1 | 4,396,284 | 1 | 4251 | 842 | 65.63996 | 3771 | 191 | 0 |
| 16 | TDR33 | L4.10 | 4,393,262 | 1 | 4235 | 855 | 65.59422 | 3740 | 199 | 1 |
| 17 | NR-123 | L4.10 | 4,414,860 | 1 | 4253 | 847 | 65.61805 | 3755 | 217 | 0 |
| 18 | H37Rv | L4.10 | 4,417,942 | 1 | 4257 | 846 | 65.617744 | 3757 | 218 | 0 |
| 19 | TMC102 | L4.10 | 4,417,942 | 1 | 4249 | 840 | 65.617744 | 3757 | 218 | 0 |
| 20 | BCCM_092 | L4.6.2.2 | 4,393,623 | 1 | 4256 | 833 | 65.60734 | 3759 | 200 | 2 |
| 21 | TDR69 | L4.6.1.2 | 4,405,078 | 1 | 4261 | 853 | 65.595955 | 3754 | 213 | 0 |
| 22 | TDR58 | L4.6.1.2 | 4,408,138 | 1 | 4266 | 856 | 65.59345 | 3757 | 214 | 0 |
| 23 | TDR51 | L4.6.1.2 | 4,408,199 | 1 | 4263 | 852 | 65.593864 | 3757 | 213 | 0 |
| 24 | BCCM_087 | L2 | 4,401,212 | 1 | 4243 | 848 | 65.62013 | 3755 | 198 | 2 |
| 25 | BCCM_3241 | L2.2.2 | 4,416,769 | 1 | 4267 | 858 | 65.643005 | 3772 | 195 | 1 |
| 26 | TDR196 | L2.2.3 | 4,409,639 | 1 | 4258 | 863 | 65.611244 | 3760 | 202 | 3 |
| 27 | TDR86 | L2.2 | 4,401,267 | 1 | 4246 | 858 | 65.61908 | 3749 | 203 | 2 |
| 28 | TDR19 | L2.2.10 | 4,407,905 | 1 | 4257 | 848 | 65.608925 | 3774 | 199 | 0 |
| 29 | Strain210 | L2.2.8 | 4,410,319 | 1 | 4285 | 862 | 65.603035 | 3786 | 197 | 1 |
| 30 | HN878 | L2.2.8 | 4,414,576 | 1 | 4277 | 860 | 65.59139 | 3781 | 201 | 0 |
| 31 | BCCM_085 | L2.2.1.1 | 4,640,384 | 1 | 4579 | 916 | 65.60871 | 3994 | 225 | 20 |
| 32 | BCCM_088 | L2.2.1 | 4,417,521 | 1 | 4271 | 851 | 65.60476 | 3775 | 202 | 1 |
| 33 | BCCM_090 | L3 | 4,439,067 | 1 | 4302 | 871 | 65.62158 | 3758 | 235 | 3 |
| 34 | BCCM_091 | L3 | 4,433,908 | 1 | 4303 | 864 | 65.61029 | 3776 | 223 | 1 |
| 35 | TDR188 | L3 | 4,431,121 | 1 | 4287 | 870 | 65.62746 | 3764 | 227 | 1 |
| 36 | TDR34 | L3.1.2.1 | 4,436,869 | 1 | 4267 | 854 | 65.63663 | 3756 | 234 | 5 |
| 37 | BCCM_089 | L3 | 4,423,147 | 1 | 4284 | 863 | 65.60865 | 3761 | 219 | 0 |
| 38 | BCCM_101 | L7 | 4,409,659 | 1 | 4274 | 880 | 65.59609 | 3741 | 206 | 5 |
| 39 | BCCM_102 | L1.1.1 | 4,419,465 | 1 | 4259 | 857 | 65.60047 | 3749 | 212 | 3 |
| 40 | TDR1 | 1.1.2 | 4,394,213 | 1 | 4237 | 863 | 65.618256 | 3729 | 209 | 1 |
| 41 | BCCM_083 | L1.1.2 | 4,421,378 | 1 | 4270 | 872 | 65.61076 | 3734 | 225 | 3 |
| 42 | TDR5 | L1.2.2 | 4,425,030 | 1 | 4268 | 869 | 65.60762 | 3737 | 226 | 2 |
| 43 | TDR146 | L1.2.2 | 4,424,317 | 1 | 4259 | 866 | 65.60866 | 3728 | 236 | 4 |
| 44 | BCCM_082 | L1.2.1 | 4,409,272 | 1 | 4275 | 860 | 65.61378 | 3755 | 215 | 5 |
| 45 | BCCM_096 | L5 | 4,421,090 | 1 | 4295 | 853 | 65.5876 | 3773 | 217 | 1 |
| 46 | BCCM_095 | L5 | 4,426,364 | 1 | 4296 | 863 | 65.59413 | 3763 | 220 | 0 |
| 47 | BCCM_097 | L5 | 4,430,669 | 1 | 4301 | 859 | 65.585945 | 3767 | 223 | 1 |
| 48 | BCCM_100 | L6 | 4,391,621 | 1 | 4244 | 850 | 65.595894 | 3755 | 177 | 1 |
| 49 | BCCM_099 | L6 | 4,387,610 | 1 | 4238 | 849 | 65.59913 | 3753 | 189 | 7 |
| 50 | BCCM_098 | L6 | 4,386,480 | 1 | 4231 | 845 | 65.59444 | 3746 | 184 | 1 |

### Phylogenetic analysis

A wgMLST maximum likelihood tree, that uses specific alleles identified through whole-genome sequencing to create a high-resolution genetic tree, was constructed based on 90,7201 amino acids in 2,891 core genes and 755 accessory genes to define the evolutionary relationships between the 50 strains (Fig. 1). All strains clustered based on their predefined lineage assignments, which were determined using Mykrobe, a bioinformatic tool that uses whole genome sequencing to identify genomic lineages and predict drug resistance [24]. Of the 50 strains, 23 (46%) belonged to Lineage 4, 9 (18%) to Lineage 2, 5 (10%) to Lineage 3, 6 (12%) to Lineage 1, 1 (2%) to Lineage 7, and 3 (6% each) to Lineages 5 and 6, respectively (Table 1). This distribution broadly mirrors global distribution of lineages where L4 accounts the majority of categorized strains, followed by L1, L2/3, L5/6, and finally L7 [25] (Fig. 1). Lineages segregated as expected with “modern” (L2, L3 and L4) lineages sharing their most recent ancestors and the “ancient” (L1, L5, 6, and L7) lineages shown as outgroups. However, inclusion of both the core and accessory gene content allowed for a more complete definition of strain relationships. The resulting tree is notable for not producing any low confidence regions unlike other similar trees [26] confirming the high quality of our sequencing platform and the Bact-Builder assemblies.

**Fig 1.**
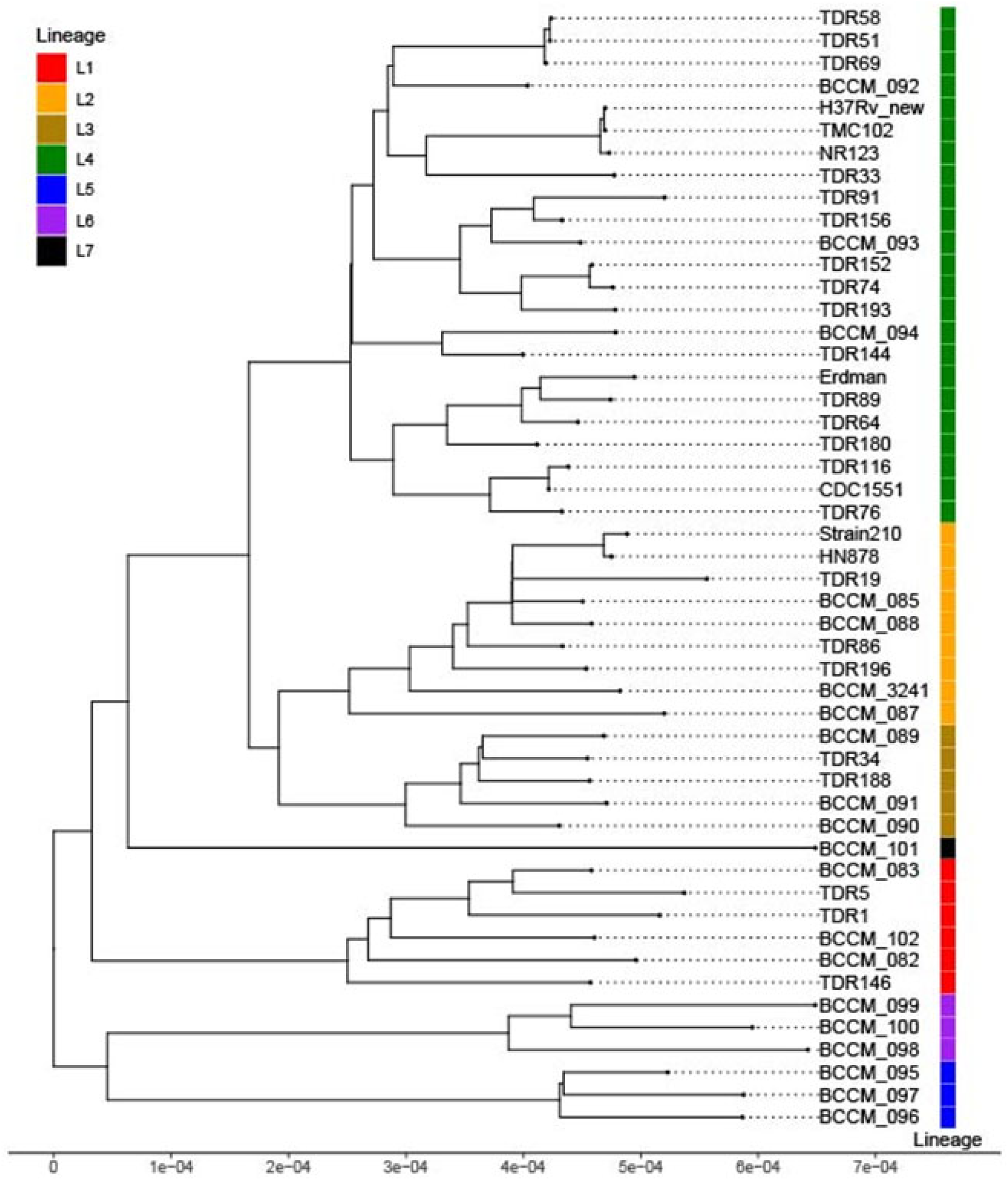
Single nucleotide polymorphism-based phylogenetic tree of all M. tuberculosis pangenome strains. A maximum likelihood phylogenetic tree was inferred using the Q.mammal+F+I subsitiution model. The x-axis represents the branch length which corresponds to the evolutionary distance as expected substitutions per site. The tree includes 50 sequences with 907,201 amino-acid sites including 2,585 parsimony informative sites. Analysis revealed expected major *M. tuberculosis* lineages and relationships among lineages. Overall distribution of strains roughly matches global lineage distribution.

### Evaluating SNPs and INDELs

A comparison of SNPs and INDELs relative to the reference H37Rv strain showed that Lineage 4 strains possessed the fewest number of SNPs and INDELs relative to H37Rv (Fig. 2A and 2B; Table 2). Lineage 1 had the highest number of SNPs and Lineage 5 had the highest number of INDELs relative to H37Rv (Fig. 2A and 2B). This finding was consistent with other studies [8, 27] and not surprising since the comparator H37Rv strain was a lineage 4 strain and. Interestingly, a comparison of intra-lineage point mutations revealed that L4 (average SNPs = 1,100) along with L1 (average SNPs = 1,182) had relatively high number of SNPs within their respective lineages compared to the other lineages (Fig. 2C and 2D; Tables S2 and S3). Moreover, L4 also harbored the highest relative number of INDELs (mean INDEL count = 966), closely followed by L1 (mean INDEL count = 811) While we observed this greater frequency of intra-lineage point mutations in L1 and L4, these differences were not found to be significant in a pairwise comparison in terms of SNPs (*p* = 0.37) or INDELs (*p* = 0.06). There were fewer INDELs on average across the lineages compared to SNPs (Table S2 and S3). The largest SNP difference was 3,300 SNPs between H37Rv (L4) and BCCM_083 (L1), and the highest INDEL difference 2,729 INDELs was between TDR 89 (L4) and BCCM_095 (L5) (Fig. 2A and 2B).

**Fig 2.**
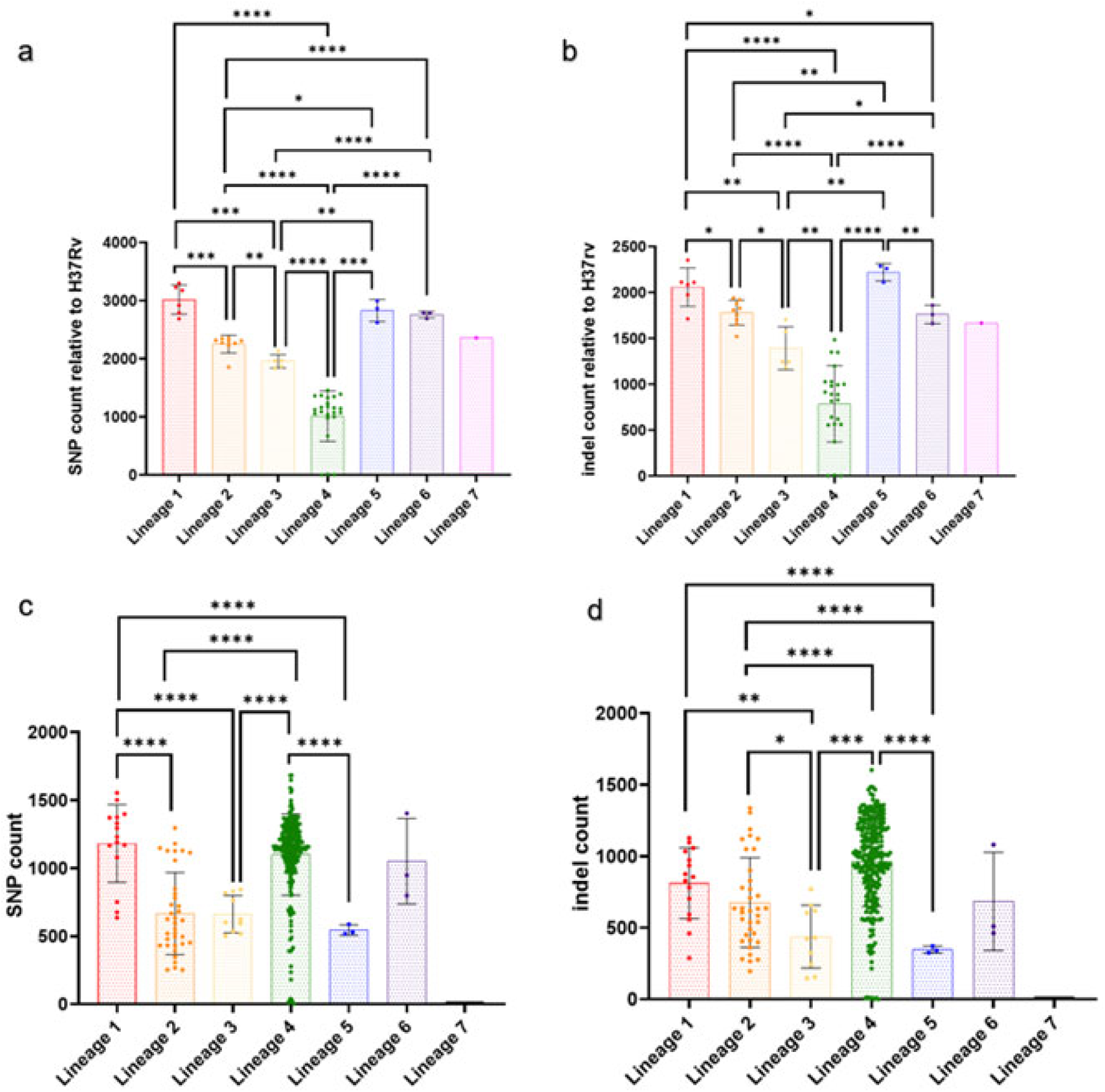
Summary of SNP and INDEL differences across all strains. a-b. Average SNP (a) and INDEL (b) values for each lineage compared to H37Rv. c-d. Average SNP (c) and INDEL (d) values for each lineage. Lineage 7 has no bar because there was only 1 L7 sample. Means + SD are depicted. Significance determined using pairwise t-tests adjusted using FDR multiple testing correction (* = P < 0.05, ** = P < 0.01, *** = P < 0.001, and **** = P < 0.0001). Nonsignificant values excluded from figure.

**Table 2.** SNP and INDEL counts of strains relative to H37Rv.

| Table 2 SNP and INDEL counts of strains relative to<br>H37Rv |  |  |  |
| --- | --- | --- | --- |
| Strain # | Strain name | SNP count | INDEL count |
| 1 | TDR76 | 1389 | 1485 |
| 2 | TDR116 | 1454 | 1351 |
| 3 | CDC1551 | 1362 | 1027 |
| 4 | TDR180 | 1348 | 1199 |
| 5 | TDR64 | 1232 | 1000 |
| 6 | Erdman | 1178 | 829 |
| 7 | TDR89 | 1152 | 891 |
| 8 | TDR144 | 1330 | 1348 |
| 9 | BCCM_094 | 1368 | 1029 |
| 10 | TDR193 | 1144 | 816 |
| 11 | TDR152 | 975 | 619 |
| 12 | TDR74 | 1001 | 642 |
| 13 | BCCM_093 | 1123 | 914 |
| 14 | TDR156 | 1073 | 888 |
| 15 | TDR91 | 1161 | 982 |
| 16 | TDR33 | 672 | 372 |
| 17 | NR-123 | 13 | 8 |
| 18 | H37Rv | 0 | 0 |
| 19 | TMC102 | 4 | 0 |
| 20 | BCCM_092 | 1042 | 995 |
| 21 | TDR69 | 1071 | 565 |
| 22 | TDR58 | 1088 | 562 |
| 23 | TDR51 | 1055 | 555 |
| 24 | BCCM_087 | 1855 | 1921 |
| 25 | BCCM_3241 | 2347 | 1653 |
| 26 | TDR196 | 2267 | 1829 |
| 27 | TDR86 | 2316 | 1703 |
| 28 | TDR19 | 2357 | 1522 |
| 29 | Strain210 | 2223 | 1808 |
| 30 | HN878 | 2276 | 1754 |
| 31 | BCCM_085 | 2311 | 1940 |
| 32 | BCCM_088 | 2302 | 1877 |
| 33 | BCCM_090 | 2134 | 1705 |
| 34 | BCCM_091 | 1964 | 1575 |
| 35 | TDR188 | 1841 | 1245 |
| 36 | TDR34 | 1866 | 1186 |
| 37 | BCCM_089 | 1968 | 1244 |
| 38 | BCCM_101 | 2359 | 1666 |
| 39 | BCCM_102 | 2921 | 2113 |
| 40 | TDR1 | 3175 | 2352 |
| 41 | BCCM_083 | 3298 | 2110 |
| 42 | TDR5 | 3227 | 2086 |
| 43 | TDR146 | 2790 | 1713 |
| 44 | BCCM_082 | 2688 | 1973 |
| 45 | BCCM_096 | 2864 | 2287 |
| 46 | BCCM_095 | 2623 | 2112 |
| 47 | BCCM_097 | 2995 | 2261 |
| 48 | BCCM_100 | 2692 | 2052 |
| 49 | BCCM_099 | 2793 | 1760 |
| 50 | BCCM_098 | 2783 | 1661 |

### Pangenome analysis

In contrast to a single genome, a pangenome takes into consideration the entire genetic repertoire of a given set of genomes that belong to a single species or a microbial clade [28]. Gene representations within a pangenome are often referred to as a set of “core” genes” that are present in all genomes within a population and “accessory” genes that occur only in a subset of the population [28–30]. However, the sequence diversity that typically develops among orthologs across a pangenome can blur the boundaries between orthologs that would commonly be thought of as identical based on synteny and high homology levels versus paralogs that may be functionally similar but not identical. Thus, a typical pangenomic analysis identifies *de novo* “gene families” (rather than individual genes) which are shared between strains based on the sequence homology of each gene against every other gene across genomes. Highly similar members of gene families can be further distributed into “gene clusters” (which will group together genes that are highly similar to one another at the amino acid sequence level) enabling the identification of "core" (e.g., groups of near-identical genes that are present in all strains) and "accessory" gene clusters (e.g., groups of near-identical genes that are absent in at least one strain). This approach completely disregards gene synteny, so similar sequences will be grouped together in the same gene cluster even if they occur in entirely different genomic contexts. While gene synteny information can help distinguish such cases and separate genes that are identical across genomes from those that are paralogous or duplicated genes (i.e., similar sequences that occur in different genomic contexts), this approach may over-split genes that have the same sequence and serve the same function if the surrounding genomic regions have undergone rearrangements, duplications, or deletions.

We performed a pangenome analysis of the 50 Mtb strains with Anvi’o, a software platform that performs comparative analyses of multiple genomes and estimates relationships based on gene content, which revealed 3,756 total gene clusters (Fig. 3A, see Data Availability). Of these, 3,377 were conserved across all strains and classified as core gene clusters. The majority of the core gene clusters (2,820) contained a single gene from each genome (i.e., single-copy core gene clusters), while 557 of core gene clusters included at least two genes from the same genome. Copy number variation across strains resulted in different frequencies of both single and multiple core gene clusters among genomes (Table 1). The remaining 379 gene clusters were classified as accessory: 291 occurred in multiple strains while 88 were strain-specific singletons. Core gene clusters represented 94.1% to 95.5% of genes in any given Mtb genome, averaging 94.7% representation across all genomes. Rarefaction curves and Heap’s law analysis (Fig. 3B) indicated that this 50-strain collection captured the majority of Mtb genetic diversity. The extremely small Heap’s law α value (0.0126) confirmed that Mtb has a highly closed pangenome with extensive genetic conservation across clades.

**Fig 3.**
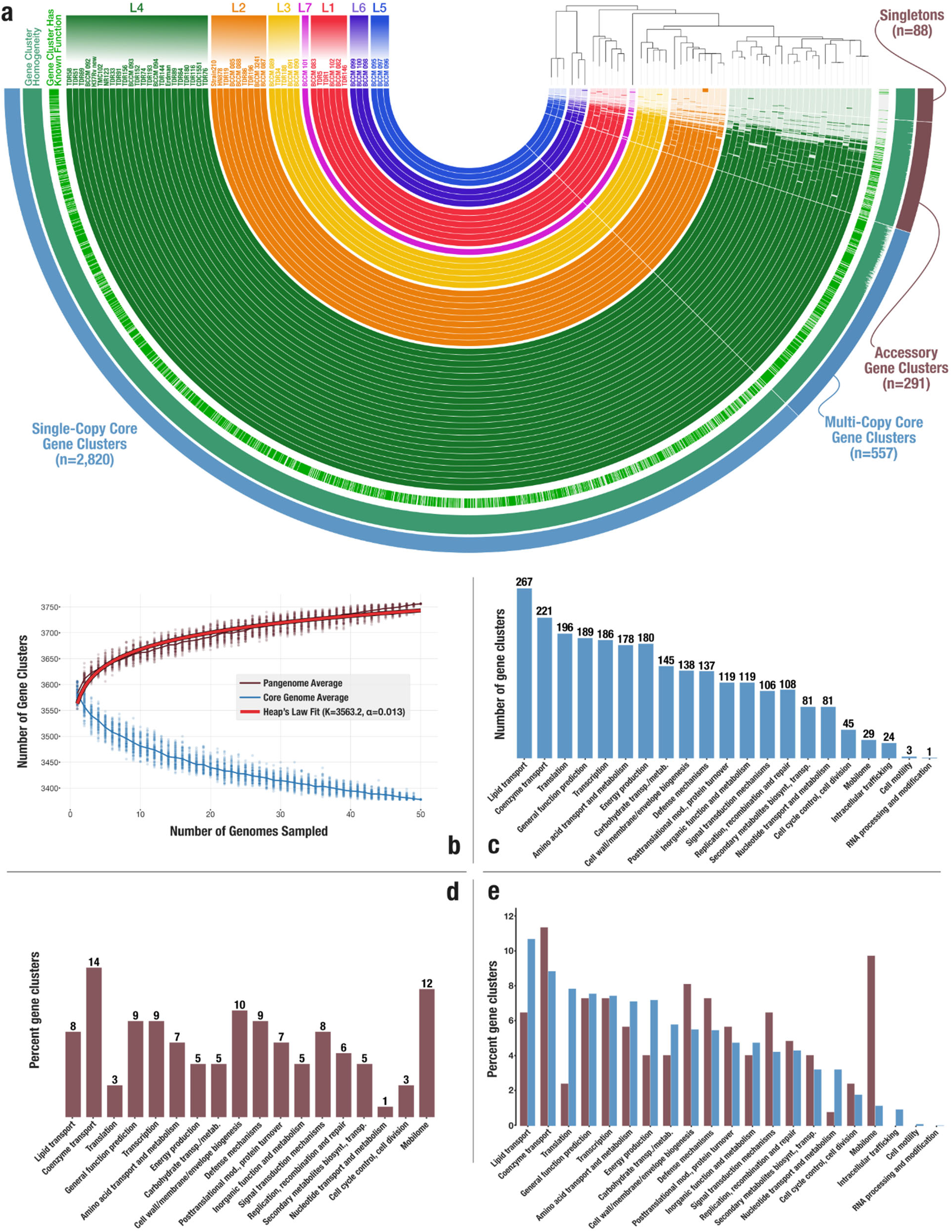
Comparative genomics of of M. tuberculosis strains. (a) The pangenomic analysis of 50 M. tuberculosis strains. Genomes are organized by the SNP tree shown in the top-right of the display and each major lineage colored. (b-d) Breakdown of functional categories of core (b) and accessory genes (c), and the comparison of the enrichment of functions in different gene pools (d). (e) The panel displays the rarefaction curves for the pangenome that summarize 100iterations along with the Heap’s Law fit.

We used the NCBI’s Clusters of Orthologous Genes (COGs) database to gain functional insights into the core and accessory gene pool. 824 of 3,377 (24.4%) core gene clusters and 253 of 379 (66.8%) accessory gene clusters had no functional annotation, suggesting that genes of unknown function comprised of a much larger fraction of the accessory gene pool. The new or previously unannotated genes in the accessory gene pool may constitute promising targets for further investigations (see Data Availability). Among those gene clusters with predicted functions, we observed 21 and 18 functional categories for core and accessory gene clusters, respectively. Most gene clusters with functional annotation in the core Mtb pangenome were involved in lipid transport (7.9% of all core gene clusters), coenzyme transport (6.5%), and translation (5.8%) (Fig. 3C, see Data Availability). In comparison, most gene clusters with functional annotation in the accessory gene pool were involved in coenzyme transport (3.7% of all accessory gene clusters), mobilome (3.2%), and cell wall/membrane biogenesis (2.6%) (Fig. 3D, see Data Availability). Side-by-side comparison of the enrichment of functional categories in core and accessory gene clusters show that mobilome genes drive the highest difference between the two pools (Fig. 3E). The higher number of mobilome genes among accessory gene clusters is most likely due to the known transposon elements that vary greatly between strains of Mtb [31, 32].

We sought to characterize the genetic makeup of multicopy gene clusters. We examined eight Mtb strains [BCCM083, BCCM088, BCCM089, BCCM093, BCCM097, BCCM099, BCCM101, and H37Rv] randomly chosen from our 50-strain set and selected 25 examples of core clusters generated by Anvi’o (Table S4). Clusters from this subset could be principally attributed to four categories of events (Table S5): 1) gene fragmentation, 2) multiple paralogs, 3) nearly identical gene duplication and 4) gene clusters caused by a combination of these features. Gene fragmentation occurred when a single open reading frame was split into two or more open reading frames by a premature stop codon followed by a secondary start codon situated downstream of the initial stop codon causing a gene “X” start-ABABABABABABABABABABABABAB-stop to be split into gene “Y” start-ABABABABABABA-stop and “Z” start-BABABABAB-stop as shown in (Fig. S1A) (Table S5, cluster 262). Here, a gene was identified in each of the eight Mtb strains examined. Seven of the eight strains had one copy of the gene, while in the eighth strain (BCCM093), the gene was split into two copies by fragmentation from a premature stop.

Multiple paralogs within one strain and/or multiple orthologs among strains was a second cause of gene clusters. In the example shown (Fig. S1B) (Table S5, cluster 145), all strains except BCCM097 contained two paralogs of the canonical Rv3269. One of the paralogs present on each genome existed as either Rv3269-1 or Rv3269-2 which differed from each other by one nonsynonymous SNP. The second paralog present on each genome existed as Rv3269-3 or Rv3269-4 which also differed from each other by one nonsynonymous SNP. In the example of H37Rv, this strain contained "Rv3269-2" & "Rv3269-3" paralogs while BCCM101 contained "Rv3269-2" & "Rv3269-4" paralogs. BCCM097 only contained one of the two paralogs.

Nearly identical gene duplications was a third cause of gene clusters. In the example shown (Fig. S1C) (Table S5, cluster 013), two strains had an additional seventh copy of the Anvi’o annotated IS285, whereas the remaining five strains had 6 copies present. Some copies were identical, others differed by only one nonsynonymous SNP and/or were of varying lengths, and other copies contained short regions that differed from this common sequence more substantially. In the case of BCCM083 and BCCM101 which had 7 copies, the additional sequence only differed by one SNP, causing us to attribute copy number variation to nearly identical gene duplication rather than the presence of paralogs as described in the previous cluster which had much more variability in sequences present. In an additional example of gene clusters caused by a combination of features. We also identified clusters containing varying copies of paralogs as well as nearly identical gene duplication.

Finally, clusters could occur due to a combination of these previously described features. In the example shown (Fig. S1D) (Table S5, cluster 022) each Mtb strain contained multiple paralogs of the *plc* (phospholipase C) gene (*plcA*, *plcB* and *plcC*) in tandem. Anvi’o annotated all the open reading frames in this cluster as *plc* while other annotation programs named these paralogs separately as *plcA*, *plcB* and *plcC*, depending on the number of copies of the gene cluster present in the strain. Gene fragmentation further expanded the size of this gene cluster, where each *plc* paralog was either present as a canonical *plc* gene sequence or as one or more of the *plc* paralogs that were split due to a premature stop codon and secondary start codon causing the cluster to include between three to five copies per genome.).

19/25 (76%) of clusters were multi-copy solely due to gene fragmentation, an additional 3/25 (12%) were due to the presence of a novel paralogs in a subset of at least one strain, 1/25 (4%) was the result of a gene duplication in one strain, and 2/25 (8%) were due to a combination of these features. In addition to the findings in our random subset that gene fragmentation increased the size of gene clusters, we would also expect to find variations in gene sizes and potentially decreases in the size of gene clusters by gene fusions created through the loss of stop codons, such as the lineage 3 Rv2044c- *pncA* gene fusion that we noted in the accessory genome (Fig. 4A-B). Although annotation artifacts could also affect the size and number of gene clusters, we did not discover any annotation artifacts in our random sample of 25 clusters.

**Fig 4.**
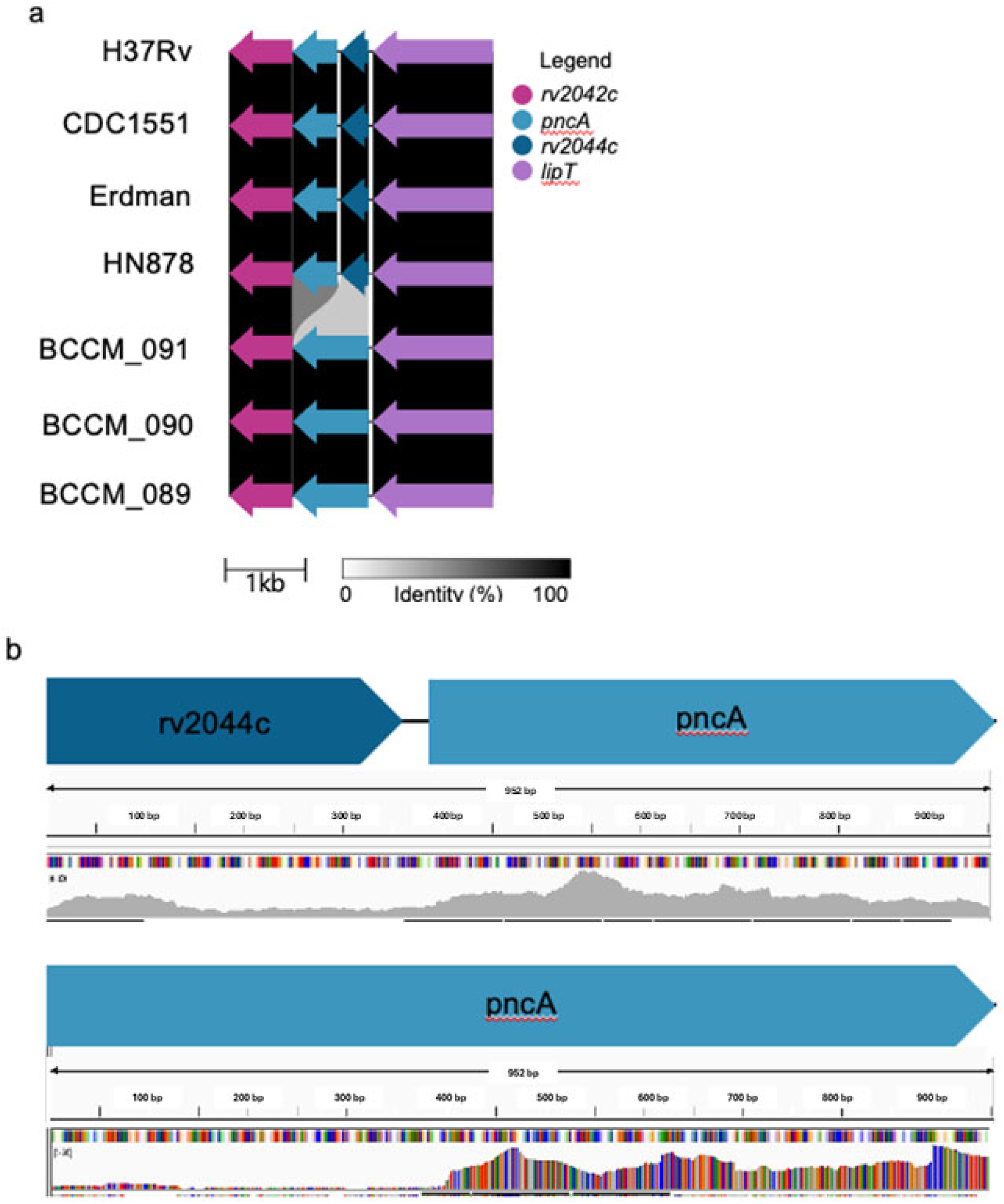
Comparison of pncA among different strains of M. tuberculosis. (a) the pncA and its surrounding genes are highly conserved across strains. A frameshift deletion rv2044c resulted in its fusion with pncA in L3 strains (BCCM_091, BCCM_090, BCCM_089). Percent similarity is shown using the gradient scale in grey. (b) Evaluation of RNAseq data (accession: PRJEB9763) aligned with minimap demonstrated that canonical pncA is still expressed and at a higher level than any fusion protein.

### A lineage 3 mutation in *pncA*

As described above, we noted that not all “new” accessory genes created by the loss of a stop codon were significantly expressed as fusion proteins. For example, a comparison of *pncA* and its surrounding genes revealed a highly conserved region (Fig. 4A), except where a frameshift deletion in the stop codon of *rv2044c*, a putative membrane protein, resulted in an out of frame gene fusion with *pncA* in all L3 strains [33]. Despite the apparent creation of this new large gene, L3 RNAseq data demonstrated that the canonical *rv2044c* and *pncA* were still separately expressed, each at high levels, compared to the new fusion RNA transcript, as can be seen by the low number of cDNA reads spanning the *rv2044c* -*pncA* junction compared to the cDNA reads aligning with the canonical open reading frame of ether gene (Fig. 4B). We assessed the functionality of PncA in strains with an *rv2044c-pncA* gene fusion by determining whether conversion of PZA to pyrazinoic acid (POA), the active form of PZA, was detectible in cultures of these strains. The two Lineage 3 strains with this gene fusions that we tested demonstrated sufficient PncA protein activity to fully hydrolyze PZA (Fig. S2). These results demonstrate that the effect of gene fusions on gene expression and bacterial phenotype should be confirmed with RNA and potentially protein or functional studies where warranted, but they also demonstrate how new genes can perhaps evolve in Mtb over time.

### Strain-specific accessory genes

We found that each of the 50 strains contained an average of 1.78 genes that were present only in that strain (Table 1). Annotation of these 89 “strain specific” accessory genes, or singletons, revealed that 29 were hypothetical genes, 3 were strain specific duplications and the remaining 37 were due to premature stop codons. All three identified duplications were found in BCCM085 and were duplications of an ATP dependent helicase, an NADH-ubiquinone oxidoreductase chain F and 3-phosphoshikimate 1-carboxyvinyltransferase respectively. The remaining accessory genes fell into the categories shown in Fig. 3D.

### Identifying hypervariable regions (HVRs)

Using PPanGGOLiN, the 50 genomes were originally divided into 468 regions of genome plasticity (RGP) comprised of genomic islands and regions that were lost across all 50 genomes. Of the identified RGPs, 19 ‘spots’ were identified that that did not have the same gene content but had similar bordering elements. Given that we were evaluating several independent strains and not ‘plastic’ changes within one strain, we renamed RGP’s as hypervariable regions (HVRs). We also developed Table S6, which simplifies the identification of each HVR (i.e. HVR 1 – 19) by listing the names and H37Rv coordinates of the conserved genes which flank each HVR. Further investigation of these spots revealed that several of them were in tandem regions that we subsequently condensed into 16 HVRs. Clinker (v0.0.25, https://github.com/gamcil/clinker) was used to visualize HVR organization [34]. Clinical strains depicted were randomly chosen as representative strains by PPanGGOLiN and commonly used reference stains (H37Rv, CDC1551, Erdman and HN878) were also included for comparison. HVR’s were broadly categorized into three groups: PE/PPE HVR’s, mobilome or transposon HVR’s, and a single lineage specific HVR.

### PE/PPE HVRs

These HVRs were characterized by differences in PE/PPE genes contained within the HVR. PE/PPE HVR’s included: HVR2, HVR5, HVR6, HVR12, HVR15, and HVR16. HVR2, bordered by *tasB* and a hypothetical protein, was caused by the insertion of transposon and mobile elements throughout the region in multiple lineages. Interestingly, although RAST annotations indicated that *PPE59* was missing in H37Rv and CDC1551, a BLAST search indicated that the sequence was present. The gene sequence for *PPE57* and *PPE58* in HVR2 however was missing in CDC1551 (Fig. S3B). HVR5, bordered by *eccC5* and *mycP5,* varied by the presence of a conserved hypothetical gene in BCCM_092 (L4) and BCCM_102 (L1) (Fig. S3E). HVR6 was bordered by *acrR* and *rv0281* (an O-methyltransferase). Variation in HVR6 was caused by the presence of a paralog of *PE_PGRS3* in several strains, the deletion of *PE_PGRS4* in HN878 and the presence of hypothetical genes scattered throughout the region in multiple lineages (Fig. S3F). HVR12 was bordered by *plcA* and *glyS*. Much of the variability in HVR12 was the result of random transposon insertions; however, we did observe duplications of *esxN.2* and *esxJ.3* and *PPE38* in TDR116 and BCCM_3241. We also observed complete or partial disruptions of *PPE38* and *PPE39* as the result of transposon insertions (Fig. S3K). HVR15 was bordered by *fadD15* and *fadD19*. HVR15 demonstrated the variability observed across PE/PPE genes. We observed variable gene sizes of *PE_PGRS55*, *PE_PGRS56* and *PE_PGRS57*. We also observed gene disruptions in *PE_PGRS53* and *PE_PGRS54* due to premature stop-codons and the presence of hypothetical genes found across multiple lineages (Fig. 5C). HVR16, bordered by *PPE54* and *rv3351c* (a putative oxidoreductase), also showed variability in PE/PPE genes. We observed variable gene sizes of *PPE54* as well as gene disruptions in *PPE54*, *PE_PGRS50*, and *PPE56* due to premature stop codons and transposon elements that were also observed across multiple lineages (Fig. S3L).

**Fig 5.**
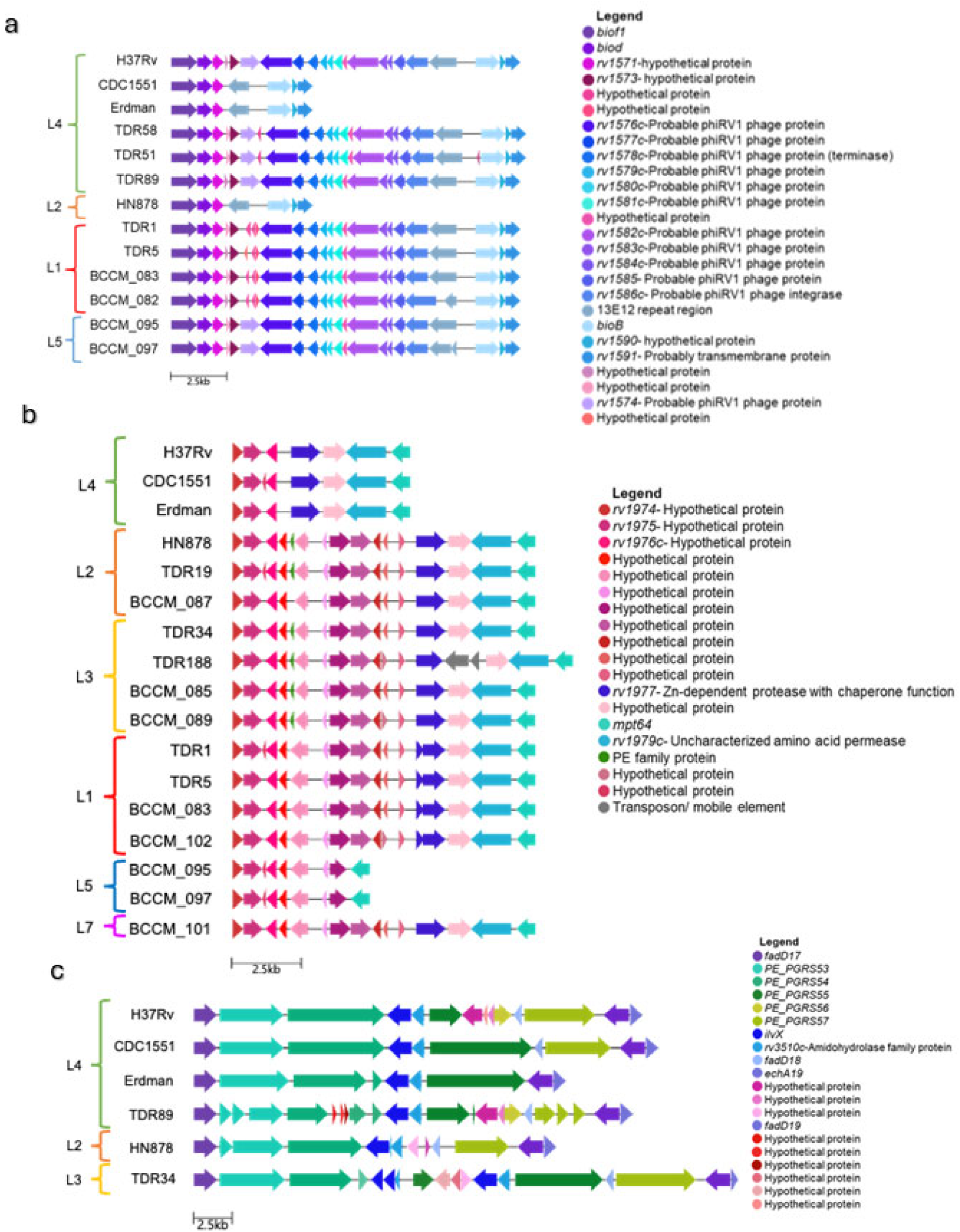
HVR regions across the M. tuberculosis Pangenome. (a-c) Schematics of (a) a mobilome or transposon HVR (b) a lineage HVR and (c) a PE/PPE HVR determined by PPanGGOLiN in representative strains and commonly used reference strains: H37Rv, CDC1551, Erdman, and HN878. Images were made using Clinker (57).

### Mobilome or transposon HVRs

These HVRs all exhibited differences in mobilome, transposon, hypothetical or phage genes contained within the HVR. HVR’s in this group are: HVR1, HVR3, HVR4, HVR7, HVR8, HVR10, HVR11, HVR13, and HVR14.

HVR1 was typified by the deletion of several phage related proteins and hypothetical proteins between border genes *bioF1* and *rv1591* in lineages 4 and 2 (Fig. 5A). The deletion was observed in CDC1551, Erdman and HN878 indicating that the deletion of this region was found in multiple lineages. HVR3, bordered by insertion sequence B9 and a hypothetical protein, was also found in multiple lineages and was caused by the deletion of several hypothetical genes in strains in Lineage 2 and 4, random transposon insertion and the deletion of *mmpl14*, a probable transmembrane transport protein, in representative strain H37Rv (Fig. S3C). HVR4, bordered by a hypothetical protein and *csm2*, was the result of random insertions of transposon elements, the deletion of several CRISPR associated proteins (*cas1*, *csm6*, *csm5*) in HN878 and the insertion of several conserved hypothetical proteins in L1, L3, L5, L6 and L7 (Fig. S3D). HVR7, bordered by *sdaA* and cystathionine beta-lyase, was largely conserved except for the presence of small hypothetical genes in certain strains, the presence of a transposon element in BCCM_091 (L3) and the deletion of *rv0071* (retron-type RNA-directed DNA polymerase), *rv0072* (an ABC-type antimicrobial peptide transporter) and Xaa-Pro dipeptidase in HN878 (Fig. S3G). The deletion was not observed in other L2 strains. HVR8, bordered by *fadD36* and *rv1203c* (conserved hypothetical protein), was also largely conserved except for small hypotheticals found in several strains in L4, 2, 3, 1 and 7 (Fig. S3H). HVR-10, bordered by *moaX* and *dacB1*, like HVR2 and HVR3 was also largely the result of transposon and mobile elements throughout the region. Several strains across all lineages also contained several small conserved hypothetical genes. Representative strains H37Rv and Erdman also contained deletions or partial deletions of *rv3324a* (Pterin-4-alpha-carbinolamine dehydratase) and a GTP 3’,8-cyclase (Fig. S3I). HVR11, bordered by *rv3015c* (methionine synthase) and *rv3026c* (Acyl-CoA:1-acyl-sn-glycerol-3- phosphateacyltransferase), was also largely the result of transposon or mobile elements randomly inserted throughout the region. We also observed the presence of several conserved hypotheticals and the deletion of *esxR* and *esxS* in Erdman (L4), TDR116 (L4), BCCM_098 (L6) and all L3 strains (Fig. S3J). HVR13, bordered by *rv1043c* (a protease-related protein) and *rv1049* (a transcriptional regulator) was largely conserved across strains, except for the presence of additional transposon elements in certain strains and the deletion of a hypothetical protein in CDC1551 (Fig. S3M). HVR14 was bordered by *rv3463* (Coenzyme F420-dependent N5,N10-methylenetetrahydromethanopterin reductase and related flavin-dependent oxidoreductases) and *ilvB2*. Several strains had a large insertion of several hypothetical or probable phage-related proteins that were observed across multiple lineages. The phage proteins were deleted in representative strains H37Rv (L4) and HN878 (L2) (Fig. S3A).

### Lineage HVR

This category of HVR had lineage specific differences in gene content. HVR9, was bordered by *rv1974* (conserved hypothetical) and *mpt64*. Unlike the other HVRs, HVR9 exhibited differences that varied in the same way by all members of one or more specific lineages. Several hypothetical proteins present in L1, 2, 3, and 7 were deleted in lineage 4. All of the hypothetical genes in HVR9 in addition to *rv1977* (a Zn-dependent protease with chaperone function), a large hypothetical gene and *rv1979c* (an uncharacterized amino acid permease) were deleted in L5 strains. In L6, the entire region was deleted except for *mpt64* and *rv1979c* (Fig. 5B).

### The PGRR, a universal reference sequence for inter-strain studies

We compiled a Pangenome Gene Reference Resource (PGRR) comprised of all genes and their paralogs to provide an efficient tool for performing isolated gene alignments and for accessing gene-level information from our pangenome reference strains. Paralogs were defined as any gene that had at least one SNP difference across the pangenome. One potential application of the PGRR is to summarize genomic variability for genes of interest across strains or lineages. For example, individual genome variability can be easily extracted, aligned, and compared across outcomes, such as drug tolerance or drug resistance. In addition, the PGRR can be used as a universal reference sequence to efficiently align DNA- or RNA-sequencing reads from non-reference sequences (such as those obtained from clinical strains) to identify strain-specific gene content or genome variability. To explore the use of the PGRR in this capacity, we first focused on *the* Mtb genome TbD1 region, a region that is found in Lineages 1, 5, 6, and 7 but which is absent in modern Mtb lineages. TbD1 includes *mmpS6* and a longer 5’ region of *mmpL6*. The PGRR contained 4 paralogs of *mmpS6* . Each of these *mmpS6* paralogs represented a different *mmpS6* sequence variant from among all of the *mmpS6* genes identified in our 50 sequenced Mtb strains (although in the case of *mmpS6*, variants were only present in lineage 1, 5, 6 and 7). Similarly, the PGRR contained 5 paralogs of the longer *mmpL6* gene. The PGRR also contained 4 paralogs of the shorter *mmpL6* gene which is present in modern strains (e.g., Strains belonging to Lineage 2, 3 and 4). We identified 658 Illumina reads in our lineage 1 BCCM082 Mtb genome that mapped to TbD1. We then mapped these TbD1 reads to the lineage 4 H37Rv genome, where as expected only 45/658 (6.8%) of these reads aligned to the H37Rv since lineage 4 does not contain TbD1. We then mapped the 568 BCCM082 TbD1 reads to our PGRR, finding that all 658 (100%) of these reads aligned to the PGRR reference. All TbD1-mapped reads that aligned to the PGRR mapped to one or more of the 4 *mmpS6* paralogs, or one or more of the 5 full length *mmpL6* gene paralogs (Fig. 6; Table S7). None of the TbD1 reads mapped to the truncated *mmpL6* paralogs.

**Fig 6.**
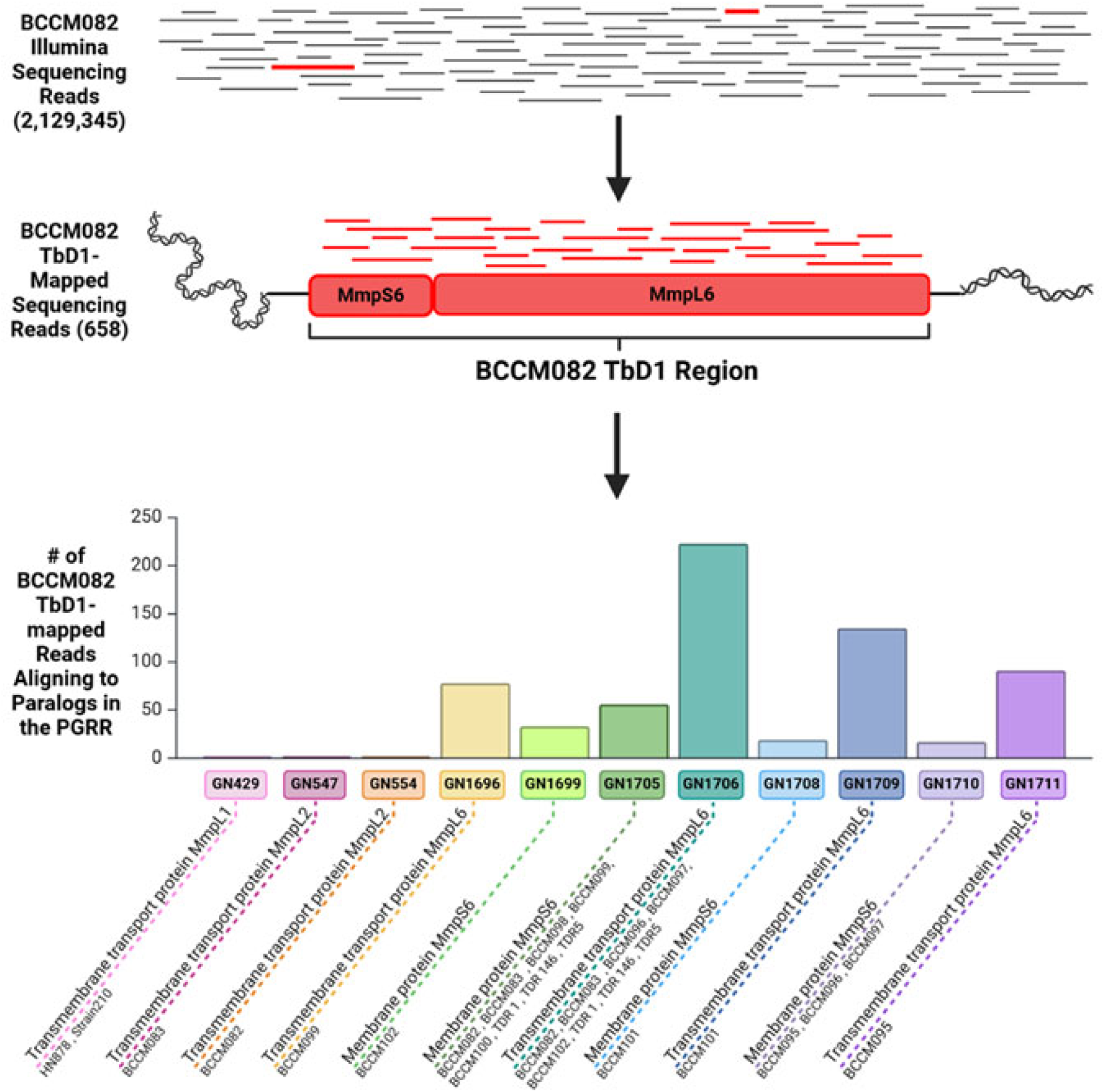
Diagram Exploring the Utility of the PGRR with a Subset of Illumina Sequencing Reads. Illumina sequencing reads from Mtb. strain BCCM082’s genomic DNA were concatenated and aligned against BCCM082’s intact TbD1 sequence. Reads mapping to this region were then aligned to the PGRR using Bowtie 2 with the “very-sensitive-local” parameter. Identifiers formatted as “GN####” represent a paralog found within the PGRR. Corresponding RAST annotations and all representative strains containing the paralog’s exact sequence from ourstudy are shown.

To evaluate the use of the PGRR to study sequencing data from whole strains, we next mapped all Illumina sequencing reads from strain BCCM082 to the complete genome of strain BCCM082 (i.e mapped to itself), H37Rv, and the PGRR using default Bowtie settings. We found that across these 2,129,345 reads, 43,964 (2.9%) aligned to the PGRR but not to H37Rv. Of the 43,964 BCCM082 reads that did not map to H37Rv, all aligned to one or more of the 17,357 paralogs of 4,294 genes present in the PGRR but not in H37Rv (Fig. S4). These genes included those with potential for drug targeting due to their roles in antimicrobial resistance, growth, and bacterial survival (Table 3). These results demonstrate that the PGRR served its expected purpose as a pangenome mapping tool to accurately match new sequence reads with paralogs already identified in the pangenome including reads that might be otherwise missed or discarded when using the H37Rv genome as the sole reference.

**Table 3:**
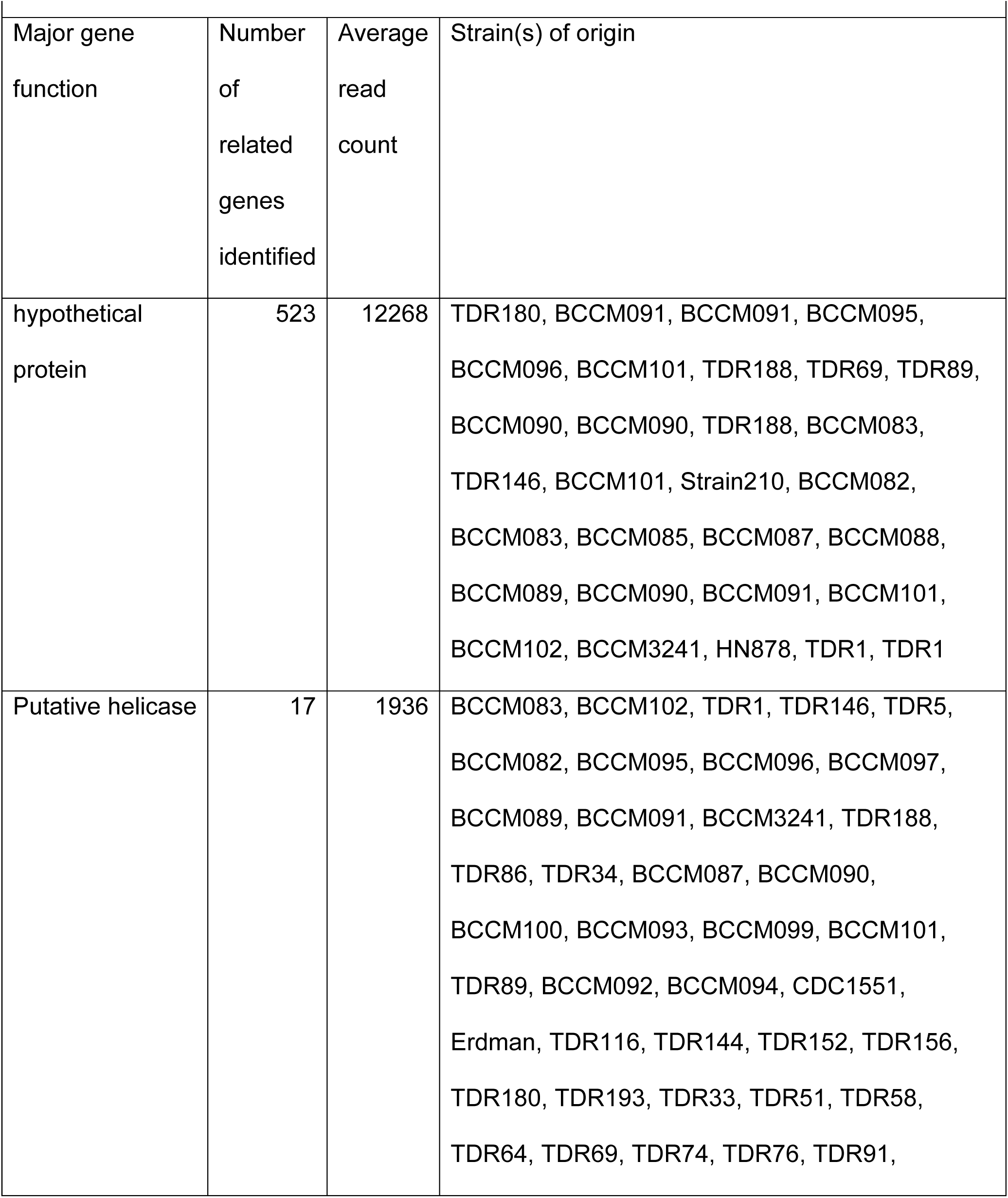
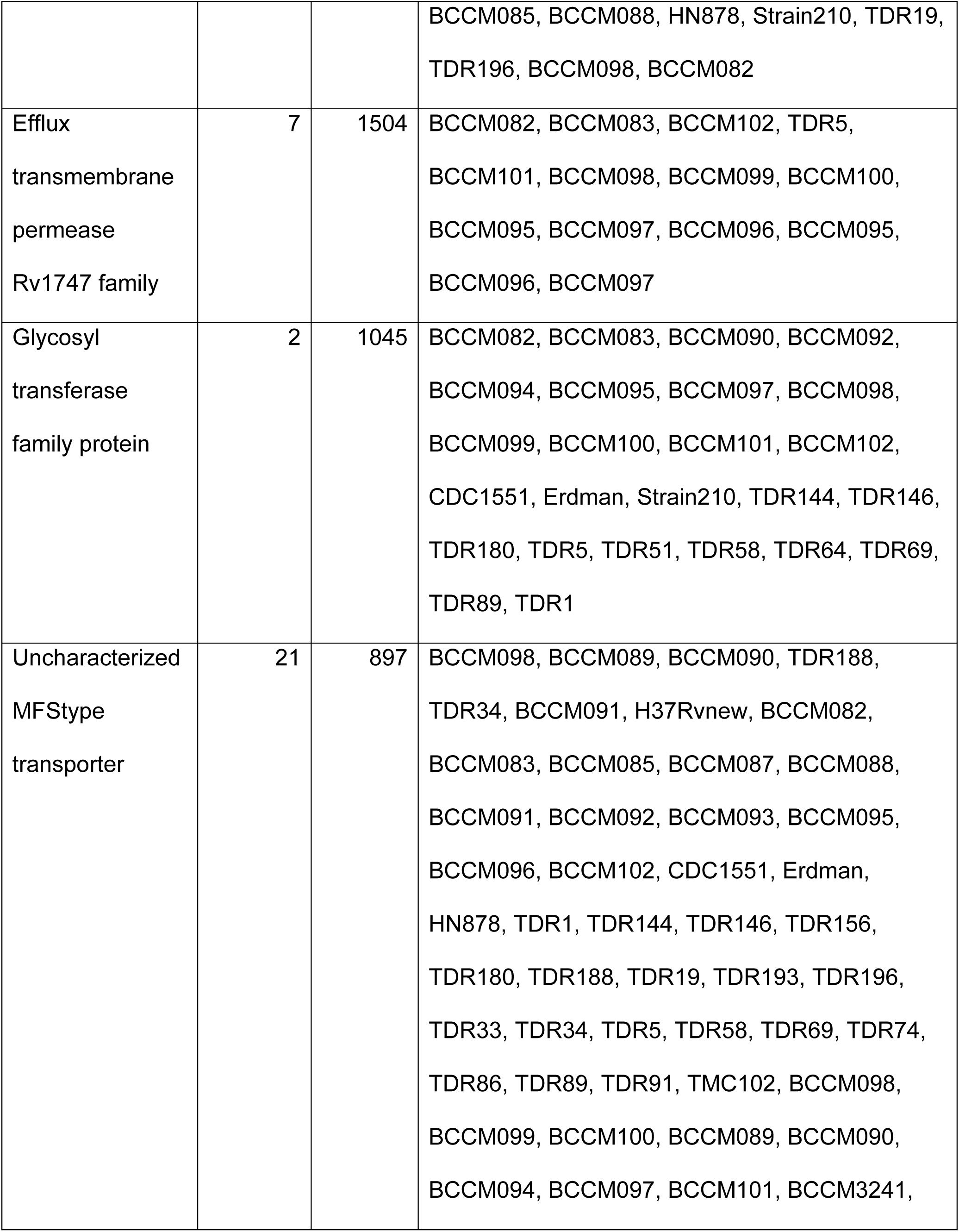
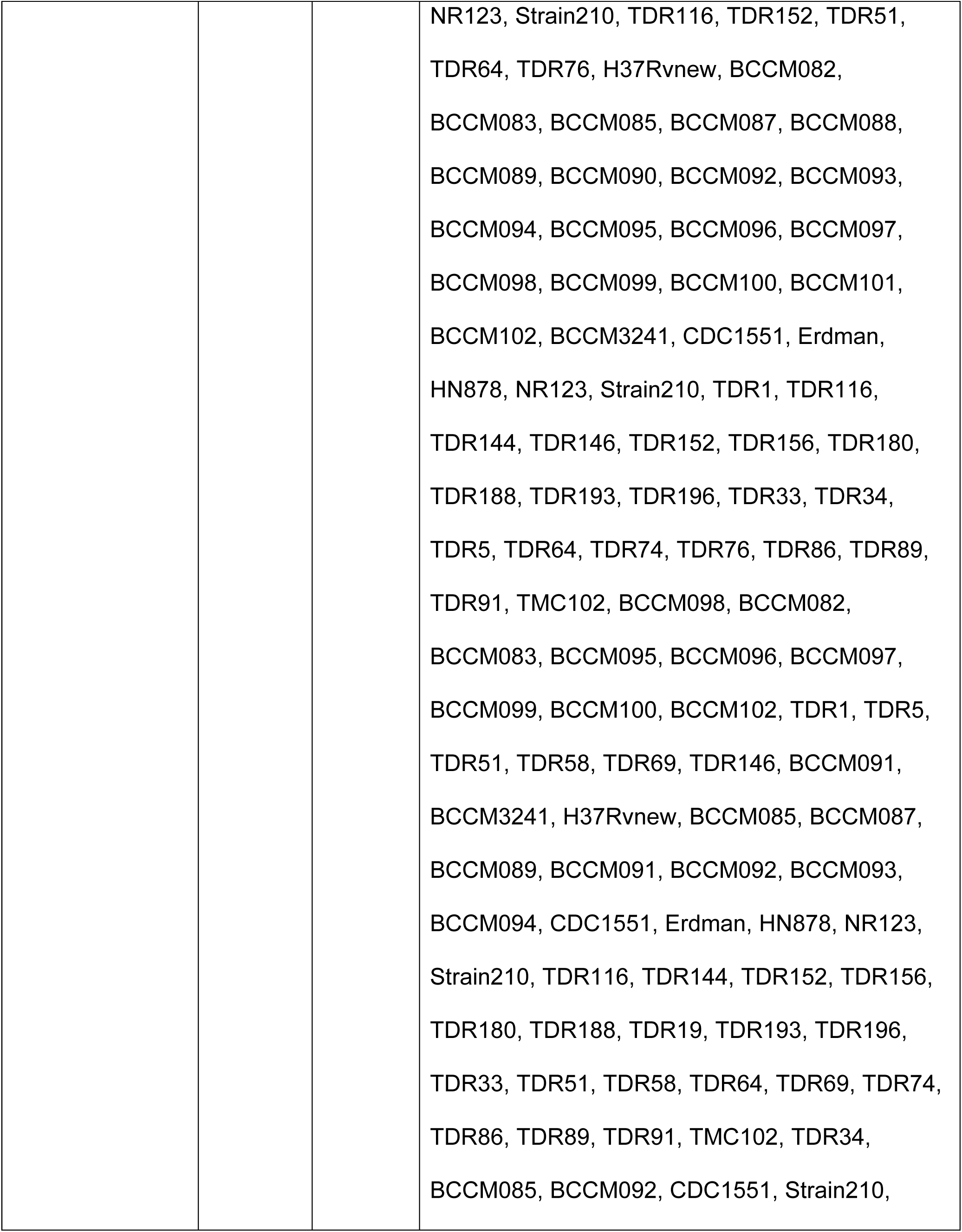
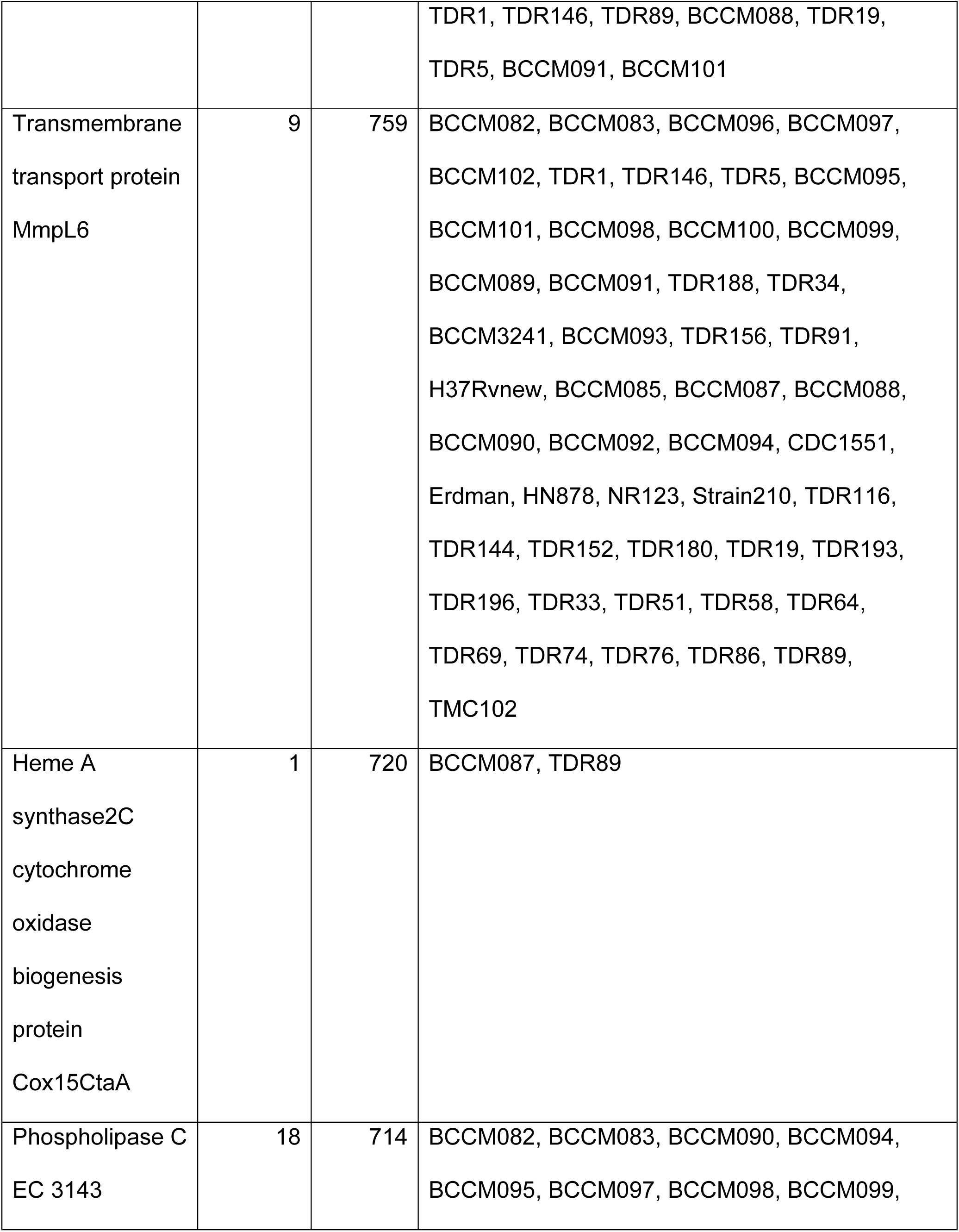
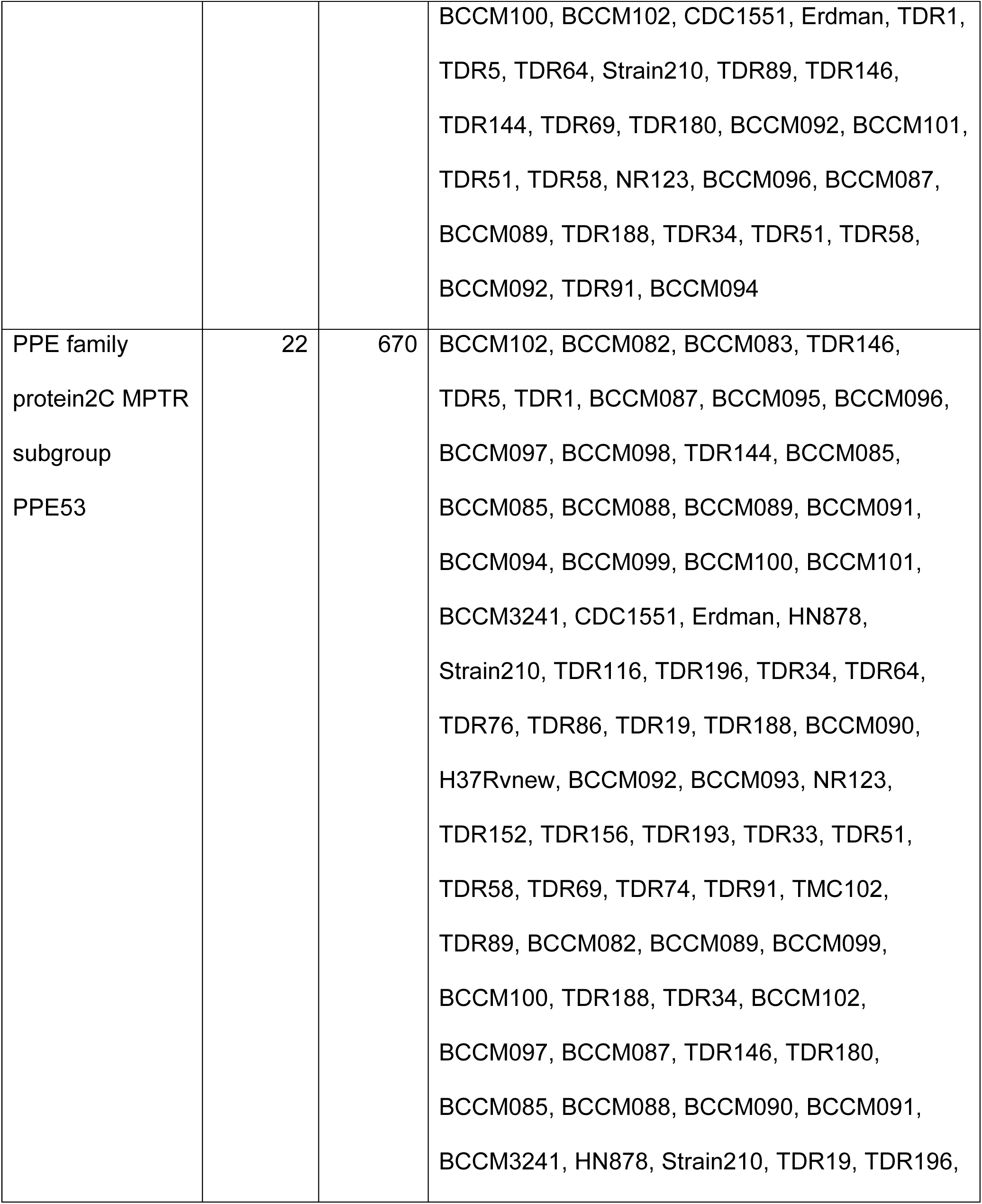
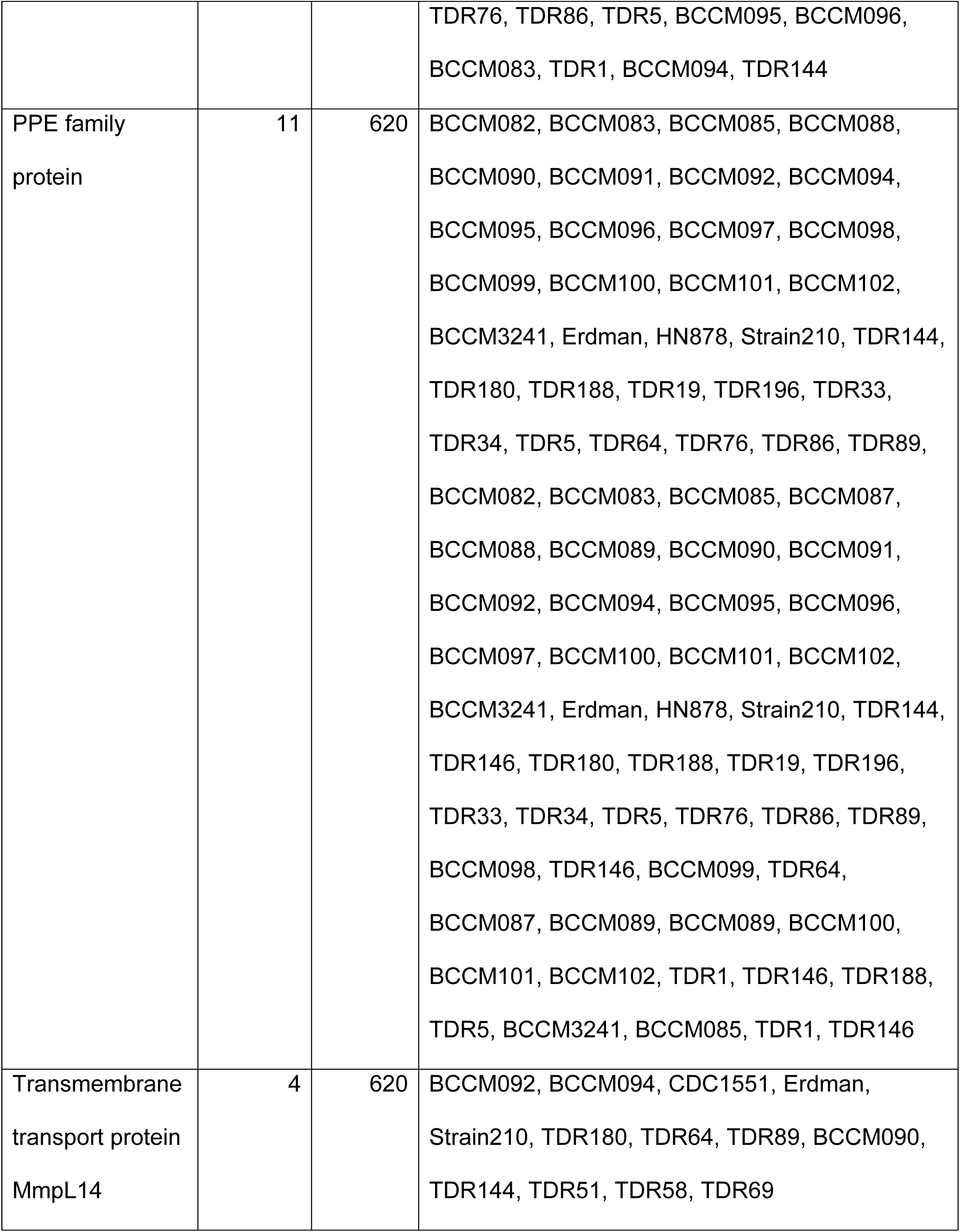
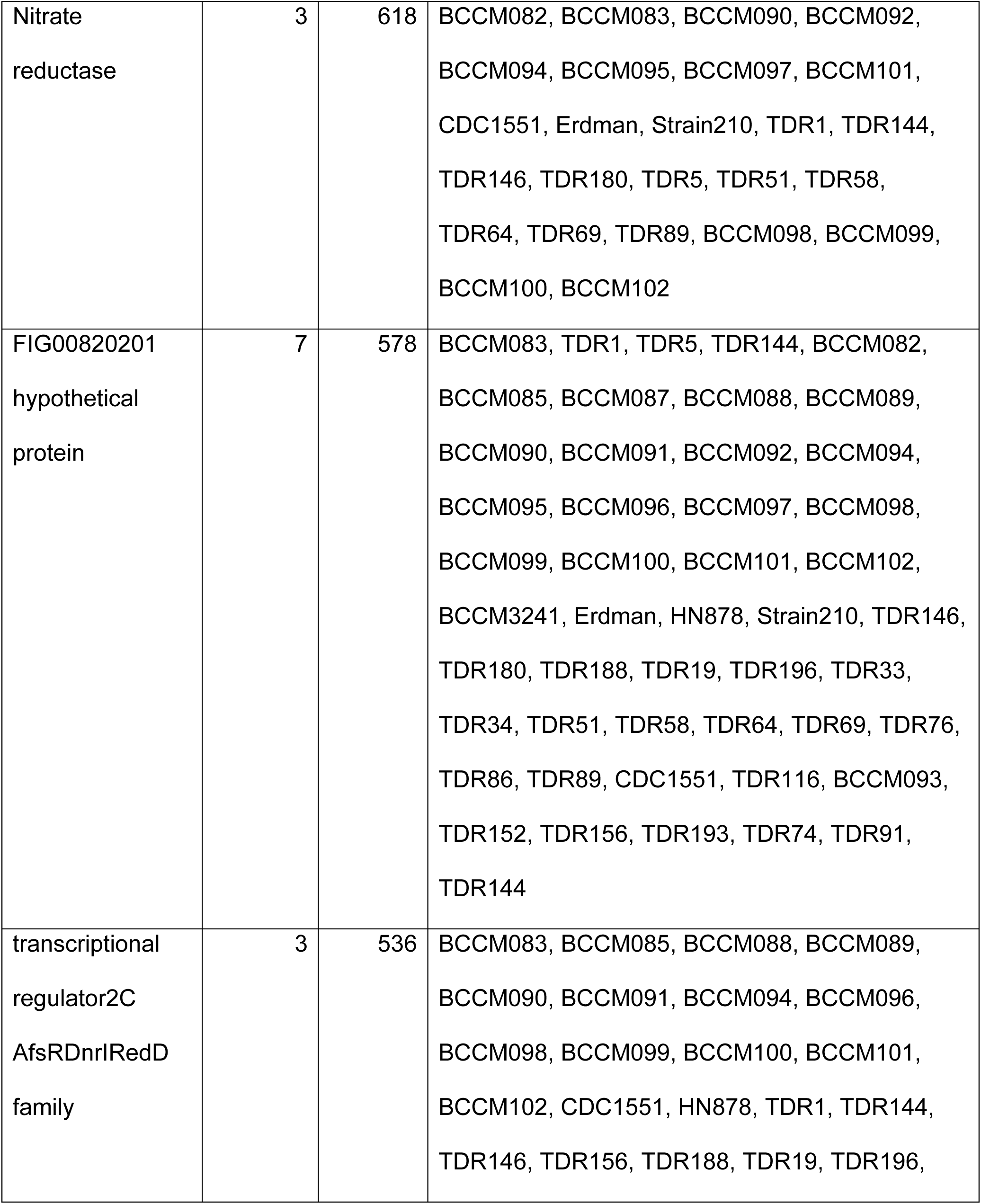
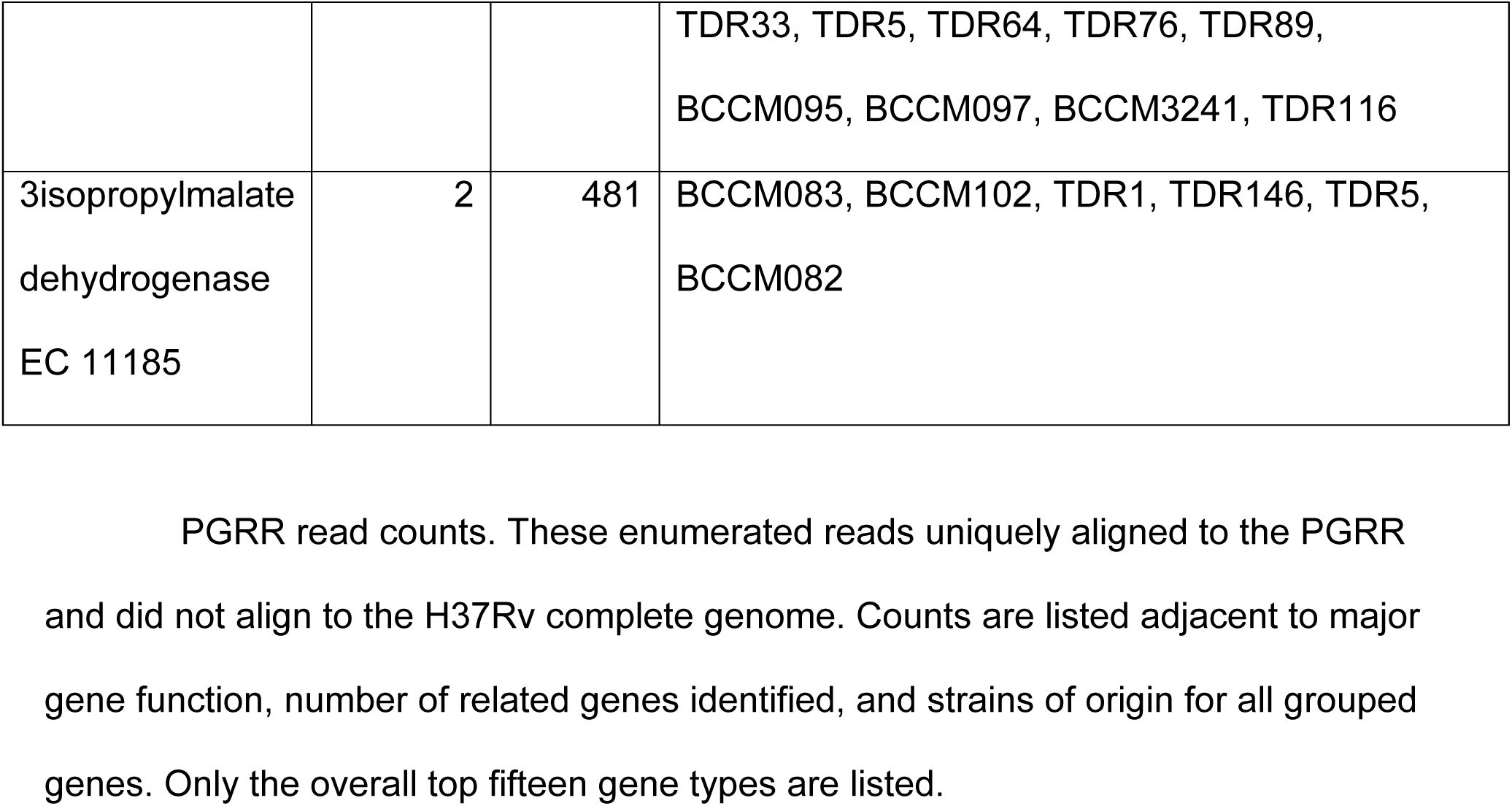
PGRR average reads counts.

## DISCUSSION

Much of Mtb research, including drug and vaccine development work, is dependent on the use of a handful of reference strains. This approach is based on the assumption that the Mtb genome is highly conserved across the entire species. We show that although the Mtb pangenome is well conserved, there exist several regions of variability that may be clinically significant. By using highly accurate, closed out genomes representing all 7 established Mtb lineages, our approach enabled a comprehensive analysis of true strain-to-strain differences across a global cohort of strains.

This work is supported by recent studies indicating that individual strains of Mtb are more diverse than previously thought [10, 18, 20, 23, 35–37]. Comparative genomic studies have also demonstrated differences in SNPs and INDELs, transposon elements and PE/PPE genes [23, 35, 38, 39]. However, most whole genome based comparative analysis studies have predominantly used H37Rv as a reference suggesting that genes not found in H37Rv were likely discarded from analysis. These studies were also predicated on the idea that the H37Rv reference genome itself is correct which we have previously demonstrated is not the case [13]. Our analysis of Mtb core and accessory genes across 50 strains supported by highly accurate sequencing and de-novo assembly methods identified a pangenome that produces variable numbers of genes within a gene cluster both within a single genome or across different strains. Our identification of multi-copy core gene clusters suggests functional relevance, as gene duplications could be driven by increasing fitness. However, gene clusters caused by gene fragmentation may in fact suggest the opposite i.e. this class of core gene may not contribute to fitness in cases where neither gene fragment is biologically useful. Our observation that some gene clusters contain both intact duplications and intact closely related paralogs as well as fragmented paralogs and gene fusions suggests further evolutionary complexity where both paralog generation and gene fragmentation within a gene cluster could both provide a fitness benefit.

We detected 16 HVR regions in our cohort. The majority of these HVRs involved variations in PE/PPE genes and gene neighborhoods. PE/PPE genes are highly repetitive and share significant homology across the entire family of genes. There are approximately 168 identified PE/PPE genes in Mtb [16, 17]. Diversity across PE/PPE genes is mostly found in the C-terminal domain of the protein. Despite the observed diversity in PE/PPE genes, we are not aware of any studies to date that have evaluated strain or lineage specific changes in PE/PPE genes or differences in sequence homology across conserved PE/PPE genes. We observed that 6/16 HVRs involved variability either in PE/PPE genes or in PE/PPE gene neighborhoods. We and others have previously described HVR-12, but a comparison across our pangenome strains revealed even more diversity than was previously known [13, 40]. We also identified duplications of PPE38, *esxN.2* and *esxJ.3* in TDR116 (L4) and BCCM_3241 (L2) (Fig. 5l). PPE38 is required for the secretion of all PE_PGRS and PPE-MPTR proteins and mutations have been linked to increased virulence [41, 42]. It is likely that there are additional differences in PE/PPE genes outside of our HVRs that will require further analysis and validation.

Six of the HVRs were the result of random transposon or mobile element insertions throughout the region. Although HVR-5 and HVR-13 were technically identified as HVRs by PPanGGOLiN, the variability in these regions were due to small annotated hypothetical genes in certain strains. Two HVRs, HVR-1 and HVR-14, included large probable prophage phiRv protein insertions in several strains. Full-length prophages are comprised of multiple components, including complete excision and integration cassette, lysis cassette, and bacteriophage structure proteins. Although the presence of variable numbers of phiRv have been previously described, their exact function in Mtb is still a mystery [17, 43].

In addition to identifying novel genes and hypervariable regions, we were able to aggregate the intergenic diversity observed across our pangenome into a single resource which we have called PGRR. This resource enables the rapid retrieval of gene sequences from a single, multiple, or all strains in the pangenome. This resource will facilitate multiple types of sequence base analyses, including the analysis of gene-level sequence variation across strains within/across gene isoforms or strains, multi-sequence alignment and phylogenetic analysis of coding sequences of TB genes, and the accurate alignment of DNA or RNA sequencing data from clinical strains. For the latter specifically, the PGRR resource will directly align sequences to tens of thousands of gene isoforms, assuring the most accurate alignment and determination of gene content within the sequenced TB strain. This application is most relevant in cases where we do not have the reference genome at hand or when the genome is not annotated. The PGRR not only enhances alignment accuracy but also accommodates the variant nature of the Mtb pangenome, reducing or eliminating false alignments and preventing sequence reads from being eliminated entirely. This tool is versatile and expansible in that it facilitates the convenient incorporation of additional genomes as they are sequenced in future amendments.

Compared to prior Mtb comparative pangenome studies [44] [45], our approach differs in several key methodological and analytical aspects. Previous pangenome analyses have often relied on large numbers of draft assemblies generated primarily from short-read sequencing, with gene presence–absence patterns inferred relative to a reference genome such as H37Rv. Other recent work has combined long- and short-read assemblies for a smaller set of isolates to investigate broad-scale structural variation, but without a formal or systematic identification of recurrent hypervariable regions (HVRs) many within or adjacent to PE/PPE genes, which we found to account for substantial diversity between strains both within and between lineages. Our strategy also enabled reference-independent calculation of lineage-specific SNP and INDEL counts, revealing patterns that differ from strictly reference-based methods, and allowed the discovery of previously undescribed duplications in virulence-associated genes such as PPE38, esxN.2, and esxJ.3. Although not comprehensive, our manual curation of a subset of core gene clusters enabled us to identify genetic drivers of gene cluster development. These findings, especially if further expanded, should provide insights into how the Mtb pangenome generates diversity in the absence of horizontal gene transfer. Furthermore, our publicly available PGRR resource provides a flexible platform for retrieving lineage- and strain-specific sequences, enabling downstream applications such as targeted variant analysis and improved clinical sequence alignment—capabilities not offered by earlier studies. These methodological and analytical advances strengthen the accuracy, resolution, and translational potential of our findings, providing a more comprehensive and reliable framework for understanding strain-specific diversity in Mtb.

While this study provides a robust and comprehensive pangenome analysis of Mtb, several limitations should be noted. Firstly, our analysis is based on 50 complete genomes, which, although diverse, may not fully capture the entire genetic diversity present in the species, particularly in underrepresented or newly discovered lineages. Additionally, while we have identified a number of hypervariable regions (HVRs), the functional relevance of these regions, particularly in relation to pathogenicity or clinical outcomes, remains an area for further investigation. Final, while our PGRR resource is a valuable tool for sequence alignment, it is limited by the availability of high-quality, annotated genome data. Continued sequencing efforts and the development of additional strain-specific resources will be essential for further refining this database and enhancing its utility in clinical and research settings.

## DATA AVAILABILITY

Whole genome sequencing data for the strains used in this study have been deposited in the NCBI sequence read archive under SRA Project number PRJNA1010209. Complete, annotated genomes for the strains in this study have been deposited in the NCBI GenBank under project number PRJNA1010209. Supplementary tables referenced in this study have been deposited to https://doi.org/10.5281/zenodo.17496644. Pangenomic analysis data from Anvio used in this study have been deposited to https://github.com/wejlab/PGRR-supplementary. The Pangenome Gene Reference Resource sequence data have been deposited to https://github.com/wejlab/PGRR.

## CODE AVAILABILITY

Zenodo: Underlying data for ‘Defining the Mycobacterium tuberculosis Pangenome and Suggestions for a New Composite Reference Sequence.’. https://doi.org/10.5281/zenodo.10426196

## ACKNOWLDEGEMENTS

Research reported in this publication was supported by the National Institute of Allergy and Infectious Diseases of the National Institutes of Health under award numbers U19AI11276 and U19AI162598 and Aubrey R. Odom was funded by T32HL125232. The content is solely the responsibility of the authors and does not necessarily represent the official views of the National Institutes of Health. The authors acknowledge the Genomics Center Rutgers New Jersey Medical School (https://research.njms.rutgers.edu/genomics/) and the Office of Advanced Research Computing (OARC) at Rutgers, The State University of New Jersey (http://oarc.rutgers.edu) for providing access to the Amarel cluster and associated research computing resources that have contributed to the results reported here.

## CONTRIBUTIONS

P.C., A.R.O., W.E.J., and D.A. designed the study; P.C., E.M.O., and C.A.G. library prepped and sequenced the samples; P.C., A.R.O., A.D.L., E.M.O., P.K., K.A.V. and E.C.F performed data analysis; P.C.,A.R.O., E.M.O., K.A.V., P.K., H.F., A.H., and A.M.E. contributed to data visualization. P.C., E.M.O., H.F., A.H., A.M.E., and D.A. wrote the manuscript. W.E.J., K.A.V., S.D.M. and P.K. edited the manuscript.

## Notes

### Competing Interest Statement

The authors have declared no competing interest.

