## Supplementary Figures for "Defining the *Mycobacterium tuberculosis* Pangenome and Suggestions for a New Composite Reference Sequence"

### Fig S1A

```
fadEd22_1      MGIALTDDHRELSGVARAFLTSQKVRWAARASLDAAGDARPPFWQNLAELGWLGLHIDER      60
fadE22_2       MGIALTDDHRELSGVARAFLTSQKVRWAARASLDAAGDARPPFWQNLAELGWLGLHIDER      60
fadE22_3       ------------------------------------------------------------      0

fadEd22_1      HGGSGYGLSELVVVIEELGRAVAPGLFVPTVIASAVVAKEGTDDQRARLLPALIDGTLTA      120
fadE22_2       HGGSGYGLSELVVVIEELGRAVAPGLFVPTVIAGAD*-----------------------      96
fadE22_3       ------------------------------------------------------------      0

fadEd22_1      GVGLDSQVQVTDGVADGEAGIVLGAGLAELLLVAAGDDVLVLERGRKGVSVDVPENFDPT      180
fadE22_2       ------------------------------------------------------------      96
fadE22_3       -------VQVTDGVADGEAGIVLGAGLAELLLVAAGDDVLVLERGRKGVSVDVPENFDPT      53

fadEd22_1      RRSGRVRLDNVRVTTDDILLGAYESALARARTLLAAEAVGGAADCVDSAVAYAKVRQQFG      240
fadE22_2       ------------------------------------------------------------      96
fadE22_3       RRSGRVRLDNVRVTTDDILLGAYESALARARTLLAAEAVGGAADCVDSAVAYAKVRQQFG      113

fadEd22_1      RTIATFQAVKHHCANMLVAAESAIAAVWDAARAAAEDEEQFRLAAAVAAALAFPAYARNA      300
fadE22_2       ------------------------------------------------------------      96
fadE22_3       RTIATFQAVKHHCANMLVAAESAIAAVWDAARAAAEDEEQFRLAAAVAAALAFPAYARNA      173

fadEd22_1      ELNIQVHGGIGFTWEHDAHLHLRRALVTVGLFGGDAPVRDVFERTAAGVTRAISLDLPAQ      360
fadE22_2       ------------------------------------------------------------      96
fadE22_3       ELNIQVHGGIGFTWEHDAHLHLRRALVTVGLFGGDAPVRDVFERTAAGVTRAISLDLPAQ      233

fadEd22_1      AEELRARIRSDAAEIAALEKDAQRDKLIETGYVMPHWPRPWGRAAGAVEQLVIEEEFSAA      420
fadE22_2       ------------------------------------------------------------      96
fadE22_3       AEELRARIRSDAAEIAALEKDAQRDKLIETGYVMPHWPRPWGRAAGAVEQLVIEEEFSAA      293

fadEd22_1      GIERPDYSITGWVILTLIQHGTPWQIERFVEKALRQQEIWCQLFSEPDAGSDAASVKTRA      480
fadE22_2       ------------------------------------------------------------      96
fadE22_3       GIERPDYSITGWVILTLIQHGTPWQIERFVEKALRQQEIWCQLFSEPDAGSDAASVKTRA      353

fadEd22_1      TRVEGGWKINGQKVWTSGAQYCARGLATVRTDPDAPKHAGITTVIIDMLAPGVEVRPLRQ      540
fadE22_2       ------------------------------------------------------------      96
fadE22_3       TRVEGGWKINGQKVWTSGAQYCARGLATVRTDPDAPKHAGITTVIIDMLAPGVEVRPLRQ      413

fadEd22_1      ITGDSEFNEVFFNDVFVPDEDVVGAPNSGWTVARATLGNERVSIGGSGSYYEAMAAKLVQ      600
fadE22_2       ------------------------------------------------------------      96
fadE22_3       ITGDSEFNEVFFNDVFVPDEDVVGAPNSGWTVARATLGNERVSIGGSGSYYEAMAAKLVQ      473

fadEd22_1      LVQRRSDAFAGAPIRVGAFLAEDHALRLLNLRRAARSVEGAGPGPEGNITKLKVAEHMIE      660
fadE22_2       ------------------------------------------------------------      96
fadE22_3       LVQRRSDAFAGAPIRVGAFLAEDHALRLLNLRRAARSVEGAGPGPEGNITKLKVAEHMIE      533

fadEd22_1      GAAIAAALWGPEIALLDGPGRVIGRTVMGARGMAIAGGTSEVTRNQIAERILGMPRDPLI      720
fadE22_2       ------------------------------------------------------------      96
fadE22_3       GAAIAAALWGPEIALLDGPGRVIGRTVMGARGMAIAGGTSEVTRNQIAERILGMPRDPLI      593

fadEd22_1      S*      721
fadE22_2       --      96
fadE22_3       S*      594
```

Fig S1B

```
plcC_1      VGSEHPVDGMTRRQFFAKAAAATTAGAFMSLAGPIIEKAYGAGPCPGHLTDIEHIVLLMQ      60
plcC_2      VVSQGAFAGMSRRAFLAKAAGAGAAAVLTDWAAPVIEKAYGAGPCSGHLTDIEHIVLCLQ      60
plcC_3      -VSQSHIGGVSRREFLAKVA-AGGAGALMSFAGPVIEKAYGAGPCSGHLTDIEHFVFFMQ      58
plcC_4      VVSQGAFAGMSRRAFLAKAAGAGAAAVLTDWAAPVIEKAYGAGPCSGHLTDIEHIVLCLQ      60
plcC_5      -VSASPLLGMSRREFLTKLTGAGAAAFLMDWAAPVIEKAYGAGPCPGHLTDIEHIVLLMQ      59
plcC_6      ------------------------------------------------------------      0
plcC_7      ---------MSRREFLTKLTGAGAAAFLMDWAAPVIEKAYGAGPCPGHLTDIEHIVLLMQ      51
plcC_8      ---------MSRREFLTKLTGAGAAAFLMDWAAPVIEKAYGAGPCPGHLTDIEHIVLLMQ      51
plcC_9      ------------------------------------------------------------      0

plcC_1      ENRSFDHYFGTLSDTRGFDDTTPPVVFAQSGWNPMTQAVDPAGVTLPYRFDTTRGPLVAG      120
plcC_2      ENRSFDHYFGTLSAVDGFDTPTP--LFQQKGWNPETQALDPTGITLPYRINTTGGPNGVG      118
plcC_3      ENRSFDHYFGTLSGTDGFNTVSP--LFQQKGWNPMTQALDATGVTMPYRFDTTRGPFLDG      116
plcC_4      ENRSFDHYFGTLSAVDGFDTPTP--LFQQKGWNPETQALDPTGITLPYRINTTGGPNGVG      118
plcC_5      ENRSFDHYFGTLSSTNGFNAASP--AFQQMGWNPMTQALDPAGVTIPFRLDTTRGPFLDG      117
plcC_6      ------------------------------------------------------------      0
plcC_7      ENRSFDHYFGTLSSTNGFNAASP--AFQQMGWNPMTQALDPAGVTIPFRLDTTRGPFLDG      109
plcC_8      ENRSFDHYFGTLSSTNGFNAASP--AFQQMGWNPMTQALDPAGVTIPFRLDTTRGPFLDG      109
plcC_9      ------------------------------------------------------------      0

plcC_1      ECVNDPDHSWIGMHNSWNGGANDNWLPAQVPFSPLQGNVPVTMGFYTRRDLPIHYLLADT      180
plcC_2      ECVNDPDHQWIAAHLSWNGGANDGWLPAQARTRS-VANTPVVMGYYARPDIPIHYLLADT      177
plcC_3      ACVNDPDHSWVAMHESWNGGVNDNWLPAQAKTRS-AAHTPTVMGYYTRQDIPIHYLLADA      175
plcC_4      ECVNDPDHQWIAAHLSWNGGANDGWLPAQARTRS-VANTPVVMGYYARPDIPIHYLLADT      177
plcC_5      ECVNDPEHQWVGMHLAWNGGANDNWLPAQATTRA-GPYVPLTMGYYTRQDIPIHYLLADT      176
plcC_6      ------------------------------------------------------------      0
plcC_7      ECVNDPEHQWVGMHLAWNGGANDNWLPAQATTRA-GPYVPLTMGYYTRQDIPIHYLLADT      168
plcC_8      ECVNDPEHQWVGMHLAWNGGANDNWLPAQATTRA-GPYVPLTMGYYTRQDIPIHYLLADT      168
plcC_9      ------------------------------------------------------------      0

plcC_1      FTVCDGYFCSLLGGTTPNRLYWMSAWIDPDGTDGGPVLIEPNIQPLQHYSWRIMPENLED      240
plcC_2      FTICDQYFSSLLGGTMPNRLYWISATVNPDGDQGGPQIVEPAIQPKLTFTWRIMPQNLSD      237
plcC_3      FTVCDRYFCSVLGPTLPNRLYWLSATIDPDGQNGGPELQSPTFQPVRRFGWRIMPQNLSD      235
plcC_4      FTICDQYFSSLLGGTMPNRLYWISATVNPDGDQGGPQIVEPAIQPKLTFTWRIMPQNLSD      237
plcC_5      FTICDGYHCSLLTGTLPNRLYWLSANIDPAGTDGGPQLVEPGFLPLQQFSWRIMPENLED      236
plcC_6      ------------------------------------------------------------      0
plcC_7      FTICDGYHCSLLTGTLPNRLYWLSANIDPAGTDGGPQLVEPGFLPLQQFSWRIMPENLED      228
plcC_8      FTICDGYHCSLLTGTLPNRLYWLSANIDPAGTDGGPQLVEPGFLPLQQFSWRIMPENLED      228
plcC_9      ------------------------------------------------------------      0

plcC_1      AGVSWKVYQNKLLGALNNTVVGYNGLVNDFKQAADPRSNLARFGISPTYPLDFAADVRNN      300
plcC_2      AGISWKVYNSKLLGGLNDTSLSRNGYVGSFKQAADPRSDLARYGIAPAYPWDFIRDVINN      297
plcC_3      AGVSWKVYRNKTLGPIS-SVLTYGSLVTSFKQSADPRSDLVRFGVAPSYPASFAADVLAN      294
plcC_4      AGISWKVYNSKLLGGLNDTSLSRNGYVGSFKQAADPRSDLARYGIAPAYPWDFIRDVINN      297
plcC_5      AGVSWKVYQNKGLGRFINTPISNNGLVQAFRQAADPRSNLARYGIAPTYPGDFAADVRAN      296
plcC_6      ------------------------------------------------------------      0
plcC_7      AGVSWKVYQNKGLGRFINTPISNNGLVQAFRQAADPRSNLARYGIAPTYPGDFAADVRAN      288
plcC_8      AGVSWKVYQNKGLGRFINTPISNNGLVQAFRQAADPRSNLARYGIAPTYPGDFAADVRAN      288
plcC_9      ------------------------------------------------------------      0

plcC_1      RLPKVSWVLPGFLLSEHPAFPVNVGAVAIVDALRILLSNPAVWEKTALIVNYDENGGFFD      360
plcC_2      TLPQVSWVVPLTVESEHPSFPVAVGAVTIVNLIRVLLRNPAVWEKTALIIAYDEHGGFFD      357
plcC_3      RLPRVSWVIPNVLESEHPAVPAAAGAFAIVNILRILLANPAVWEKTALIVSYDENGGFFD      354
plcC_4      TLPQVSWVVPLTVESEHPSFPVAVGAVTIVNLIRVLLRNPAVWEKTALIIAYDEHGGFFD      357
plcC_5      RLPKVSWLVPNILQSEHPALPVALGAVSMVTALRILLSNPAVWEKTALIVSYDENGGFFD      356
plcC_6      -----------------------------------------VWEKTALIVSYDENGGFFD      19
plcC_7      RLPKVSWLVPNILQSEHPALPVALGAVSMVTALRILLSNPAVWEKTALIVSYDENGGFFD      348
plcC_8      RLPKVSWLVPNILQSEHPALPVALGAVSMVTALRILLSNPAVWEKTALIVSYDENGGFFD      348
plcC_9      ------------------------------------------------------------      0

plcC_1      HVVPPTPP--PGTPGEFVT-VPDIDSVPGSGGIRGPIGLGFRVPCLVISPYSRGPLMVHD      417
plcC_2      HVTPLTAP--EGTPGEWIPNSVDIDKVDGSGGIRGPIGLGFRVPCFVISPYSRGGLMVHD      415
plcC_3      HVVPATAP--AGTPGEYVT-VPDIDQVPGSGGIRGPIGLGFRVPCFVISPYSRGPQMVHD      411
plcC_4      HVTPLTAP--EGTPCEWIPNSVDIDKVDGSGGIRGPIGLGFRVPCFVISPYSRGGLMVHD      415
plcC_5      HVTPPTAP--PGTPGEFVT-VPNIDAVPGSGGIRGPLGLGFRVPCIVISPYSRGPLMVSD      413
plcC_6      HVVPATAP--AGTPGEYVT-VPDIDQVPGSGGIRGPIGLGFRVPCFVISPYSRGPQMVHD      76
plcC_7      HVTPPTAP--PGTPGEFVT-VPNIDAVPGSGGIRGPLGLGFRVPCIVISPYSRGPLMVSD      405
plcC_8      HVTPPTARHRPGHPANSSR-CPTSTQY------PGPVAFVVRSVWVFAFPA--------L      393
plcC_9      --------------------------------------------------------MVSD      4

plcC_1      TFDHTSTLKLIRARFGVPVPNLTAWRDATVGDMTSTFNFAAPPNPSKPNLDHPRLNALPK      477
plcC_2      RFDHTSQLQLIGKRFGVPVPNLTPWRASVTGDMTSAFNFAAPPDPSPPNLDHPV-RQLPK      474
plcC_3      TFDHTSQLRLLETRFGVPVPNLTAWRRSVTGDMTSTFNFAVPPNSSWPNLDYPGLHALST      471
plcC_4      RFDHTSQLQLIGKRFGVPVPNLTPWRASVTGDMTSAFNFAAPPDPSPPNLDHPV-RQLPK      474
plcC_5      TFDHTSQLKLIRARFGVPVPNMTAWRDGVVGDMTSAFNFAAPPNSTRPNLSHPLLGALPK      473
plcC_6      TFDHTSQLRLLETRFGVPVPNLTAWRRSVTGDMTSTFNFAVPPNSSWPNLDYPGLHALST      136
plcC_7      TFDHTSQLKLIRARFGVPVPNMTAWRDGVVGDMTSAFNFATPPNSTRPNLSHPLLGALPK      465
plcC_8      SFRRTAAAR*--------------------------------------------------      402
plcC_9      TFDHTSQLKLIRARFGVPVPNMTAWRDGVVGDMTSAFNFATPPNSTRPNLSHPLLGALPK      64
             * :*:  :

plcC_1      LPQCVPNAVLGTVTK--TAIPYRVPFPQSMPTQETAPTRGIPSGLC*-     521
plcC_2      VAKCVPNVVLGFLN---EGLPYRVPYPQTTPVQESGPARPIPSGIC*-     517
plcC_3      VPQCVPNAALGTIN---RGIPYRVPDPQIMPTQETTPTRGIPSGPC*-     514
plcC_4      VAKCVPNVVLGFLN---EGLPYRVPYPQTTPVQESGPARPIPSGIC*-     517
plcC_5      LPQCIPNVVLGTTDGALPSIPYRVPYPQVMPTQETTPVRGTPSGLCS*     520
plcC_6      VPQCVPNAALGTIN---RGIPYRVPDPQIMPTQETTPTRGIPSGPC*-     179
plcC_7      LPQCIPNVVLGTTDGALPSIPYRVPYPQVMPTQETTPVRGTPSGLCS*     512
plcC_8      ------------------------------------------------     402
plcC_9      LPQCIPNVVLGTTDGALPSIPYRVPYPQVMPTQETTPVRGTPSGLCS*     111
```

Fig S1C

```
IS258_1      MTSSHLIDTEQLLADQLAQASPDLLRGLLSTFIAALMGAEADALCGAGYRERSDERSNQR      60
IS258_2      MTSSHLIDAEQLLADQLAQASPDLLRGLLSTFIAALMGAEADALCGAGYRERSDERSNQR      60
IS258_3      MTSSHLIDAEQLLADQLAQASPDLLRGLLSTFIAALMGAEADALCGAGYRERSDERSNQR      60
IS258_4      MTSSHLIDTEQLLADQLAQASPDLLRGLLSTFIAALMGAEADALCGAGYRERSDERSNQR      60
IS258_5      MTSSHLIDAEQLLADQLAQASPDLLRGLLSTFIAALMGAEADALCGAGYRERSDERSNQR      60
IS258_6      MTSSHLIDAEQLLADQLAQASPDLLRGLLSTFIAALMGAEADALCGAGYRERSDERSNQR      60
             ********:***************************************************

IS258_1      NGYRHRDFDTRAATIDVAIPKLRQGSYFPDWLLQRRKRAERALTSVVATCYLLGVSTRRM      120
IS258_2      NGYRHRDFDTRAATIDVAIPKLRQGSYFPDWLLQRRKRAERALTSVVATCYLLGVSTRRM      120
IS258_3      NGYRHRDFDTRAATIDVAIPKLRQGSYFPDWLLQRRKRAERALTSVVATCYLLGVSTRQM      120
IS258_4      NGYRHRDFDTRAATIDVAIPKLRQGSYFPDWLLQRRKRAERALTSVVATCYLLGVSTRRM      120
IS258_5      NGYRHRDFDTRAATIDVAIPKLRQGSYFPDWLLQRRKRAERALTSVVATCYLLGVSTRRM      120
IS258_6      NGYRHRDFDTRAATIDVAIPKLRQGSYFPDWLLQRRKRAERALTSVVATCYLLGVSTRRM      120
             **********************************************************:*

IS258_1      ERLVETLGVTKLSKSQVSIMAKELDEAVEAFRTRPLDAGPYTFLAADALVLKVREAGRVV      180
IS258_2      ERLVETLGVTKLSKSQVSIMAKELDEAVEAFRTRPLDAGPYTFLAADALVLKVREAGRVV      180
IS258_3      ERLVETLGVTKLSKSQVSIMAKELDEAVEAFRTRPLDAGPYTFLAADALVLKVREAGRVV      180
IS258_4      ERLVETLGVTKLSKSQVSIMAKELDEAVEAFRTRPLDAGPYTFLAADALVLKVREAGRVV      180
IS258_5      ERLVETLGVTKLSKSQVSIMAKELDEAVEAFRTRPLDAGPYTFLAADALVLKVREAGRVV      180
IS258_6      ERLVETLGVTKLSKSQVSIMAKELDEAVEAFRTRPLDAGPYTFLAADALVLKVREAGRVV      180
             ************************************************************

IS258_1      GVHTLIATGVNAEGYREILGIQVTSAEDGAGWLAFFRDLVARGLSGVALVTSDAHAGLVA      240
IS258_2      GVHTLIATGVNAEGYREILGIQVTSAEDGAGWLAFFRDLVARGLSGVALVTSDAHAGLVA      240
IS258_3      GVHTLIATGVNAEGYREILGIQVTSAEDGAGWLAFFRDLVARGLSGVALVTSDAHAGLVA      240
IS258_4      GVHTLIATGVNAEGYREILGIQVTSAEDGAGWLAFFRDLVARGLSGVALVTSDAHAGLVA      240
IS258_5      GVHTLIATGVNAEGYREILGIQVTSAEDGAGWLAFFRDLVA-------------------      221
IS258_6      GVHTLIATGVNAEGYREILGIQVTSAEDGAGWLAFFRDLVARGLSGVALVTSDAHAGLVA      240
             *****************************************

IS258_1      AIGATLPAAAWQRCRTHYAANHGRHNA---------------------------------      267
IS258_2      AIGATLPAAAWQRCRTHYAANHGRHNA---------------------------------      267
IS258_3      AIGATLPAAAWQRCRTHYAANLMAATPKPSWPWVRTLLHSIYDQPDAESVVAQYDRVLDA      300
IS258_4      AIGATLPAAAWQRCRTHYAANLMAATPKPSWPWVRTLLHSIYDQPDAESVVAQYDRVLDA      300
IS258_5      -----LPAAAWQRCRTHYAANLMAATPKPSWPWVRTLLHSIYDQPDAESVVAQYDRVLDA      276
IS258_6      AIGATLPAAAWQRCRTHYAANLMAATPKPSWPWVRTLLHSIYDQPDAESVVAQYDRVLDA      300
                  ****************    .

IS258_1      ------------------------------------------------------------      267
IS258_2      ------------------------------------------------------------      267
IS258_3      LTDKLPAVAEHLDTARTDLLAFTAFPKQIWRQIWSNNPQERLNREVRRRTDVVGIFPDRA      360
IS258_4      LTDKLPAVAEHLDTARTDLLAFTAFPKQIWRQIWSNNPQERLNREVRRRTDVVGIFPDRA      360
IS258_5      LTDKLPAVAEHLDTARTDLLAFTAFPKQIWRQIWSNNPQERLNREVRRRTDVVGIFPDRA      336
IS258_6      LTDKLPAVAEHLDTARTDLLAFTAFPKKIWRQIWSNNPQERLNREVRRRTDVVGIFPDRA      360

IS258_1      -------------------------------------------------------    267
IS258_2      -------------------------------------------------------    267
IS258_3      SIIRLVGAVLAEQHDEWIEGRRYLGLEVLTRARAALTSTEEPAKQQTTNTPALTT    415
IS258_4      SIIRLVGAVLAEQHDEWIEGRRYLGLEVLTRARAALTSTEEPAKQQTTNTPALTT    415
IS258_5      SIIRLVGAVLAEQHDEWIEGRRYLGLEVLTRARAALTSTEEPAKQQTTNTPALTT    391
IS258_6      SIIRLVGAVLAEQHDEWIEGRRYLGLEVLTRARAALTSTEEPAKQQTTNTPALTT    415
```

### Fig S1D

```
Rv3269_1      MVTHELLVKAAGAVLTGLVGVSAYETVRKALGTAPIRRASVTVMEWGLRGTRRAEAAAES      60
Rv3269_2      MVTHELLVKAAGAVLTGLVGVSAYETLRKALGTAPIRRASVTVMEWGLRGTRRAEAAAES      60
Rv3269_3      MAIQVFLAKATTTVITGLAGVTAYEILKKAAAKAPLRQTAVSAAALGLRGTRKAEEAAES      60
Rv3269_4      MAIQVFLAKATTTVITGLAGVTAYEILKKAAAKAPLRQTAVSAAALGLRGTRKAEEAAES      60
              *. : :*.**: :*:***.**:*** ::** ..**:*:::*:.   ******:** ****

Rv3269_1      ARLTVADVVAEARGRIGEEAPLPAGARVDE--- 90
Rv3269_2      ARLTVADVVAEARGRIGEEAPLPAGARVDE--- 90
Rv3269_3      ARLKVADVMAEARERIGEESPTPAISDLHDHDH 93
Rv3269_4      ARLKVADVMAEARERIGEESPTPAISDLHDLDH 93
              ***.****:**** *****:* ** : :.:
```

**Fig S1.** Alignment of all unique amino acid sequences for the different causes of copy number variation observed within the core genome; including (A) a gene split due to premature stop codon, (B) a combination of ambiguous annotations and a gene split, (C) a gene duplication, or (D) the presence of a paralog.

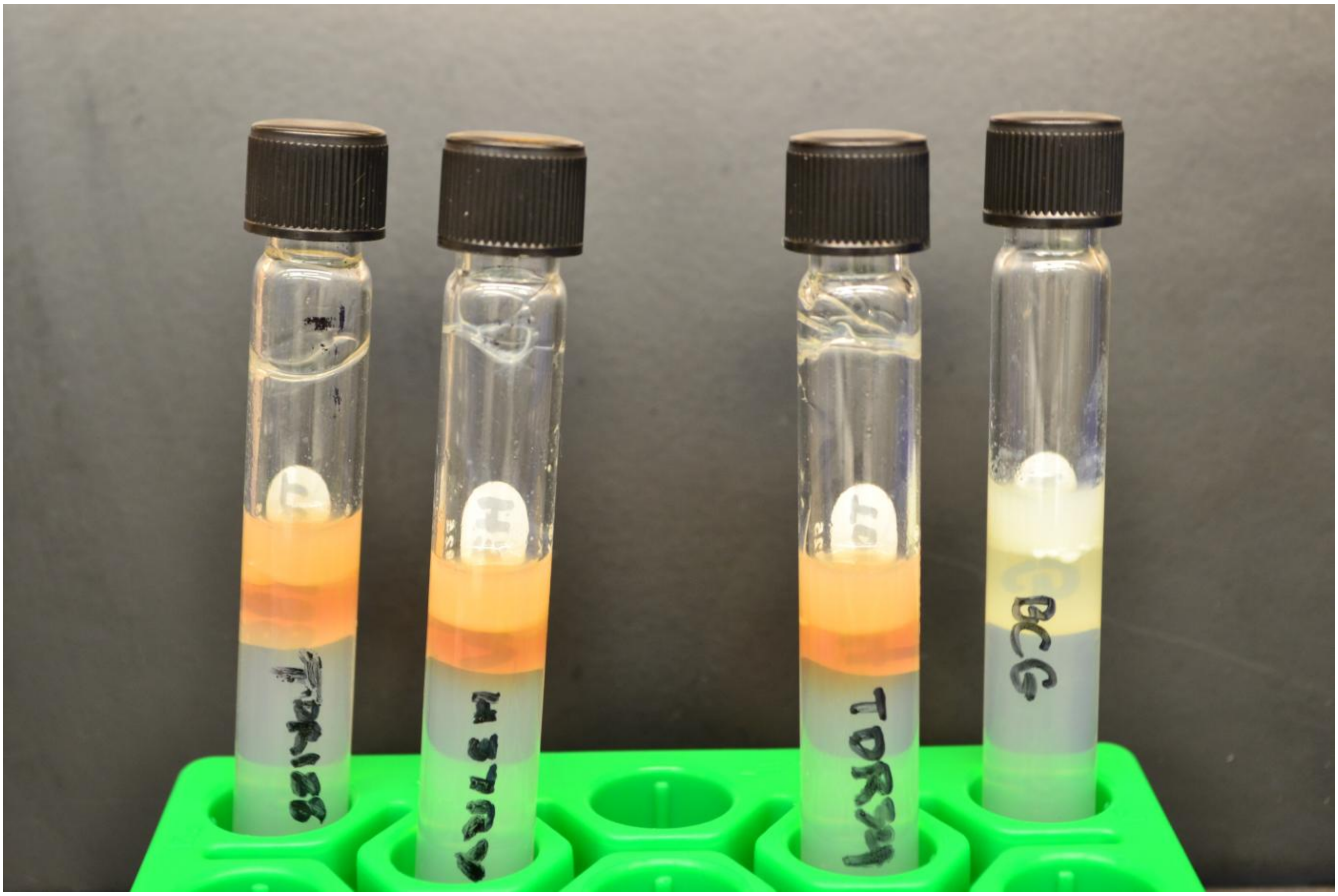


Fig. S2. Detection of pyrazinamidase activity in two Lineage 3 strains (TDR188 & TDR34), H37Rv, and BCG which is known to not have pyrazinamidase activity. Cultures were treated with PZA and the color orange demarcates the detection of the pyrazinamidase product, POA, in culture.

Fig. S3A

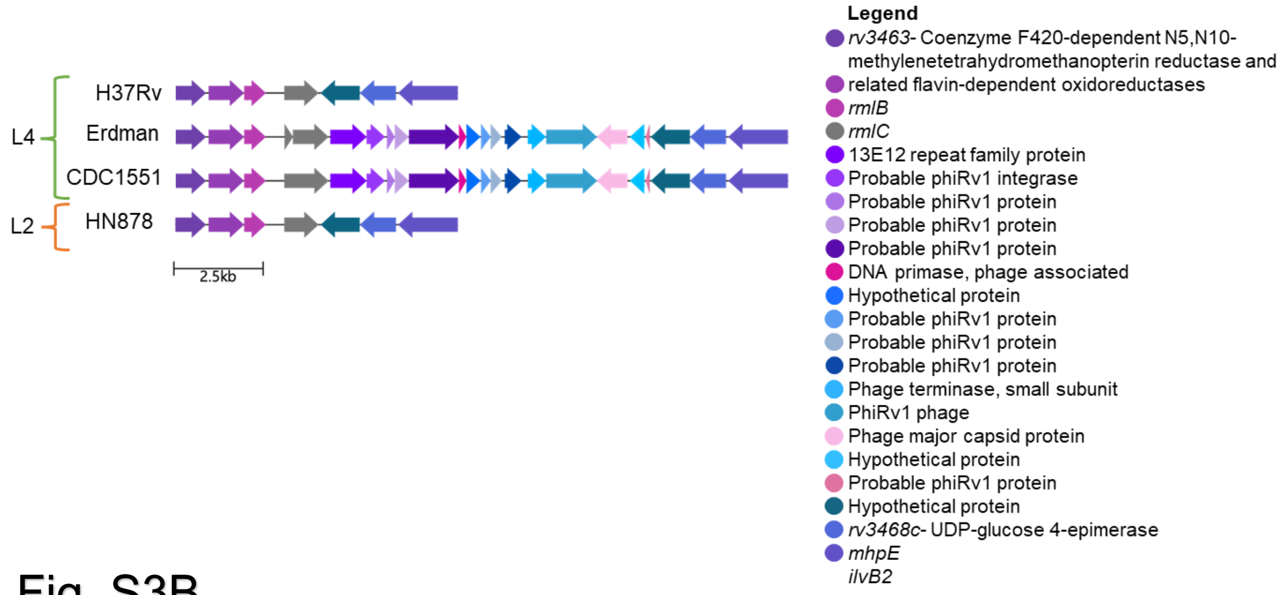


Fig. S3B

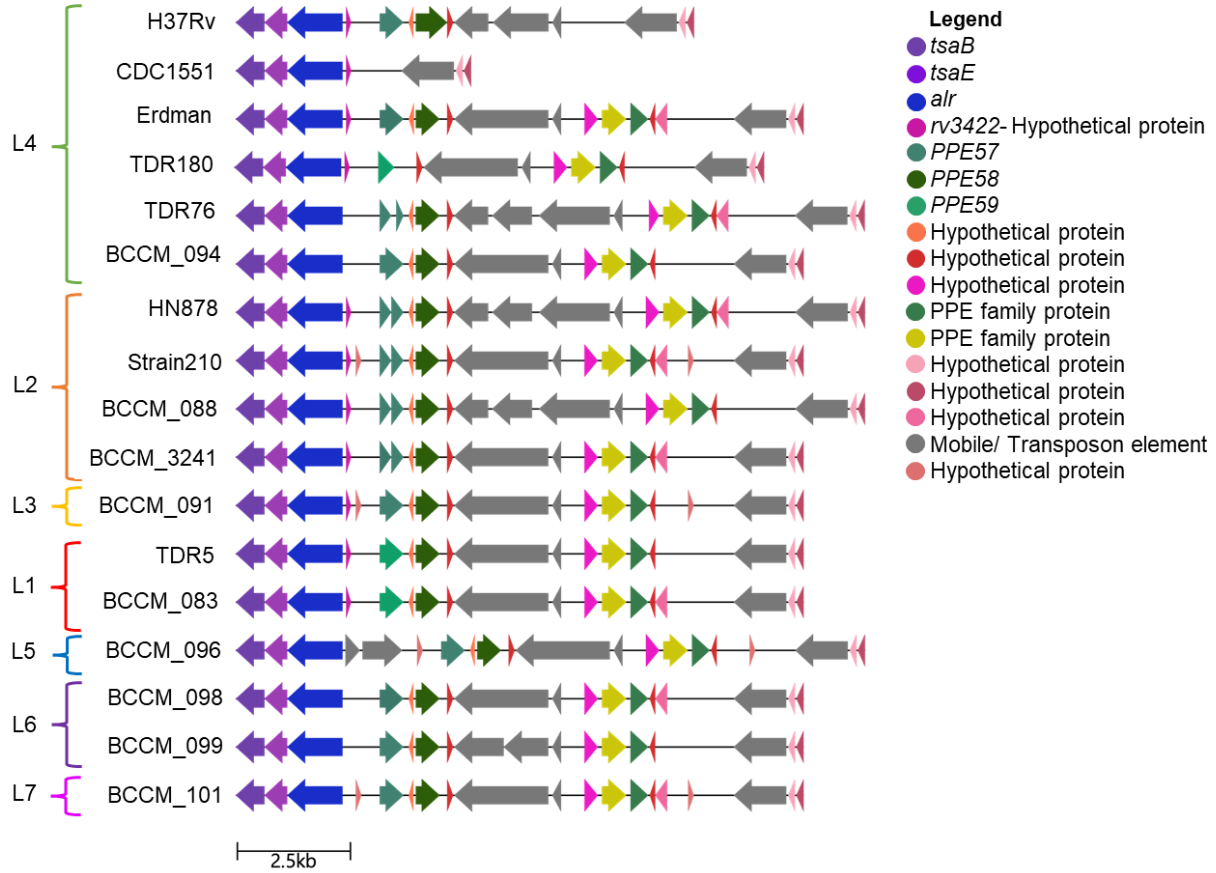

### Fig. S3C

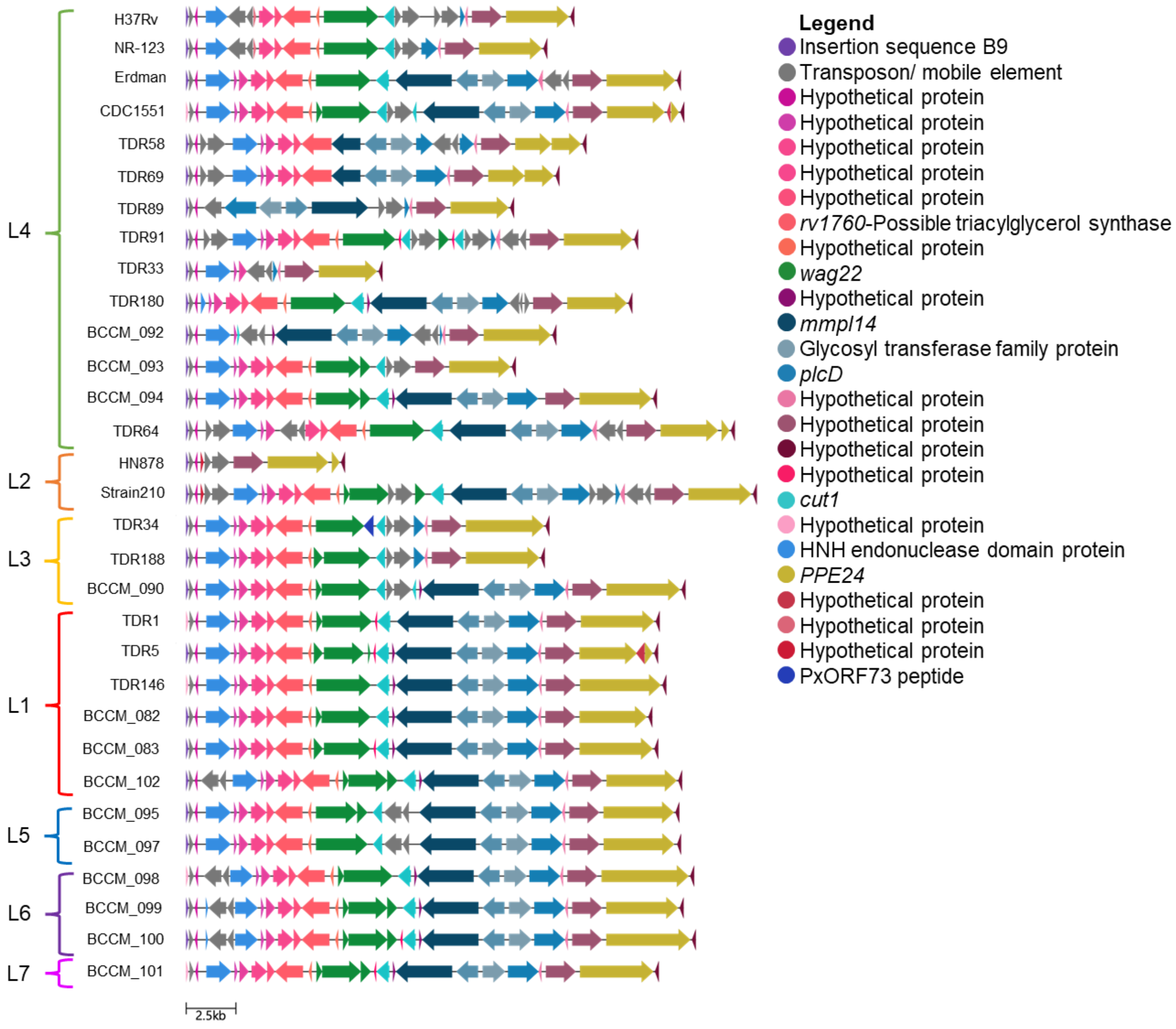

H37Rv
NR-123
Erdman
CDC1551
TDR58
TDR69
TDR89
TDR91
TDR33
TDR180
BCCM_092
BCCM_093
BCCM_094
TDR64
HN878
Strain210
TDR34
TDR188
BCCM_090
TDR1
TDR5
TDR146
BCCM_082
BCCM_083
BCCM_102
BCCM_095
BCCM_097
BCCM_098
BCCM_099
BCCM_100
BCCM_101
L4
L2
L3
L1
L5
L6
L7
2.5kb
Legend
Insertion sequence B9
Transposon/ mobile element
Hypothetical protein
Hypothetical protein
Hypothetical protein
Hypothetical protein
Hypothetical protein
rv1760-Possible triacylglycerol synthase
Hypothetical protein
wag22
Hypothetical protein
mmpl14
Glycosyl transferase family protein
plcD
Hypothetical protein
Hypothetical protein
Hypothetical protein
Hypothetical protein
cut1
Hypothetical protein
HNH endonuclease domain protein
PPE24
Hypothetical protein
Hypothetical protein
Hypothetical protein
PxORF73 peptide

Fig. S3D

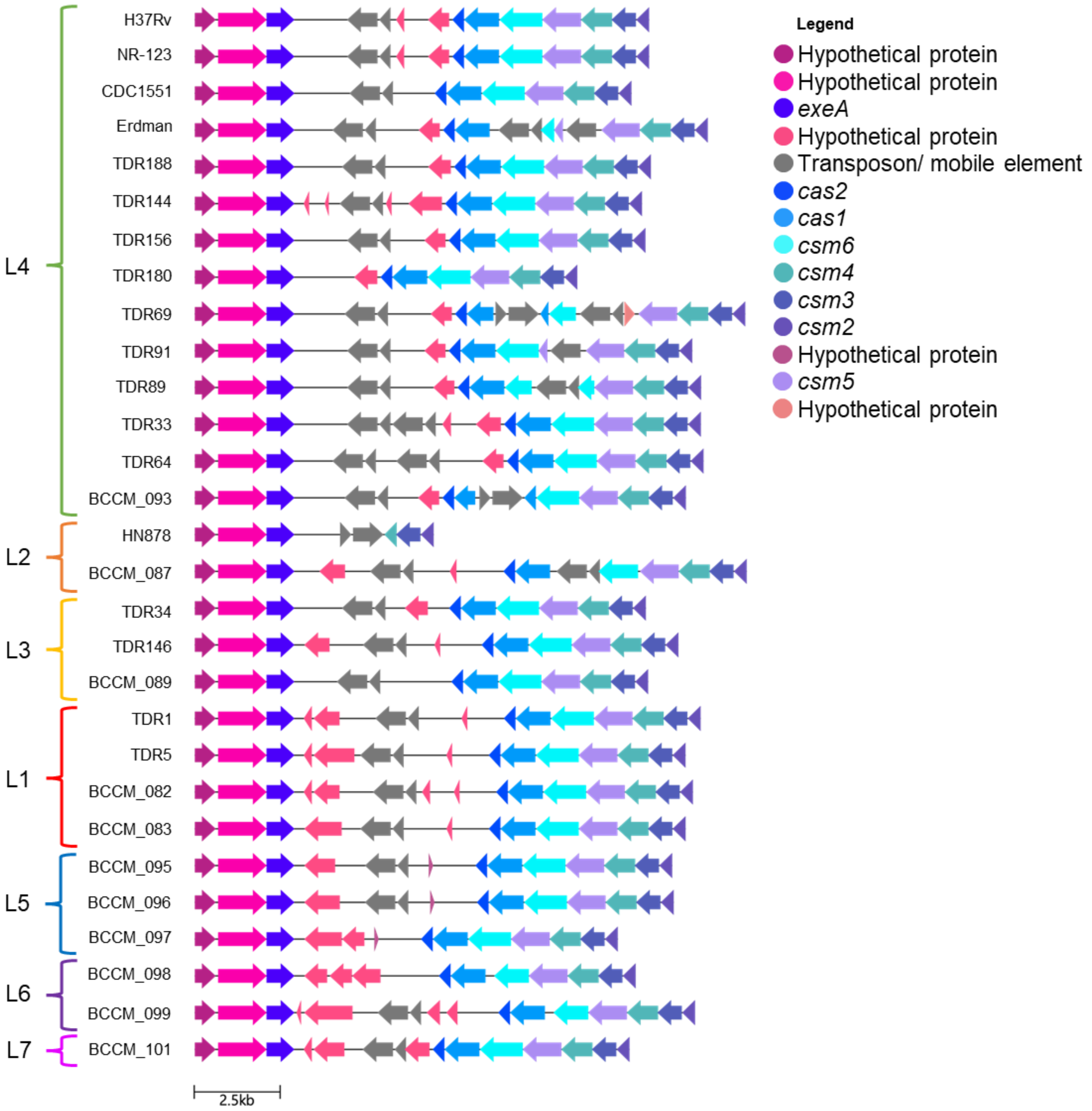
L4
H37Rv
NR-123
CDC1551
Erdman
TDR188
TDR144
TDR156
TDR180
TDR69
TDR91
TDR89
TDR33
TDR64
BCCM_093
L2
HN878
BCCM_087
L3
TDR34
TDR146
BCCM_089
L1
TDR1
TDR5
BCCM_082
BCCM_083
L5
BCCM_095
BCCM_096
BCCM_097
L6
BCCM_098
BCCM_099
L7
BCCM_101
2.5kb
Legend
Hypothetical protein
Hypothetical protein
exeA
Hypothetical protein
Transposon/ mobile element
cas2
cas1
csm6
csm4
csm3
csm2
Hypothetical protein
csm5
Hypothetical protein

### Fig. S3E

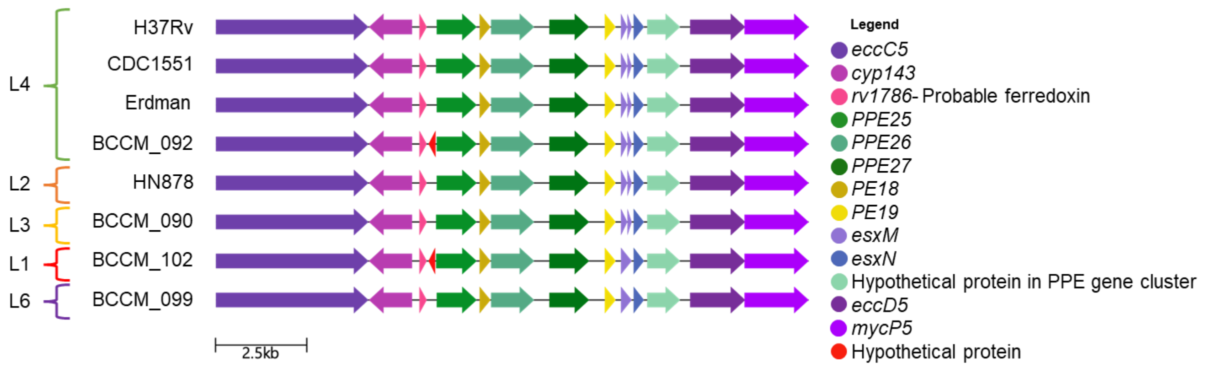


### Fig. S3F

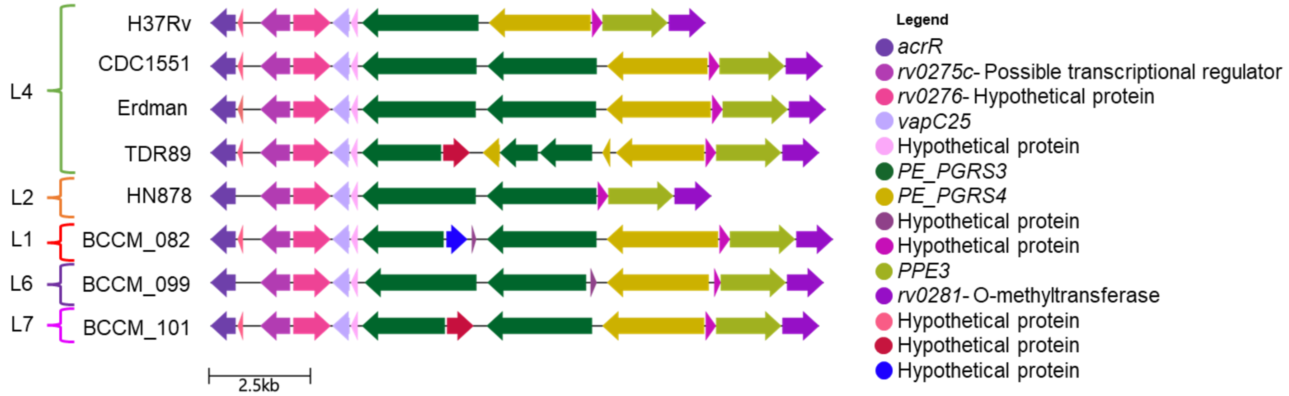


### Fig S3G

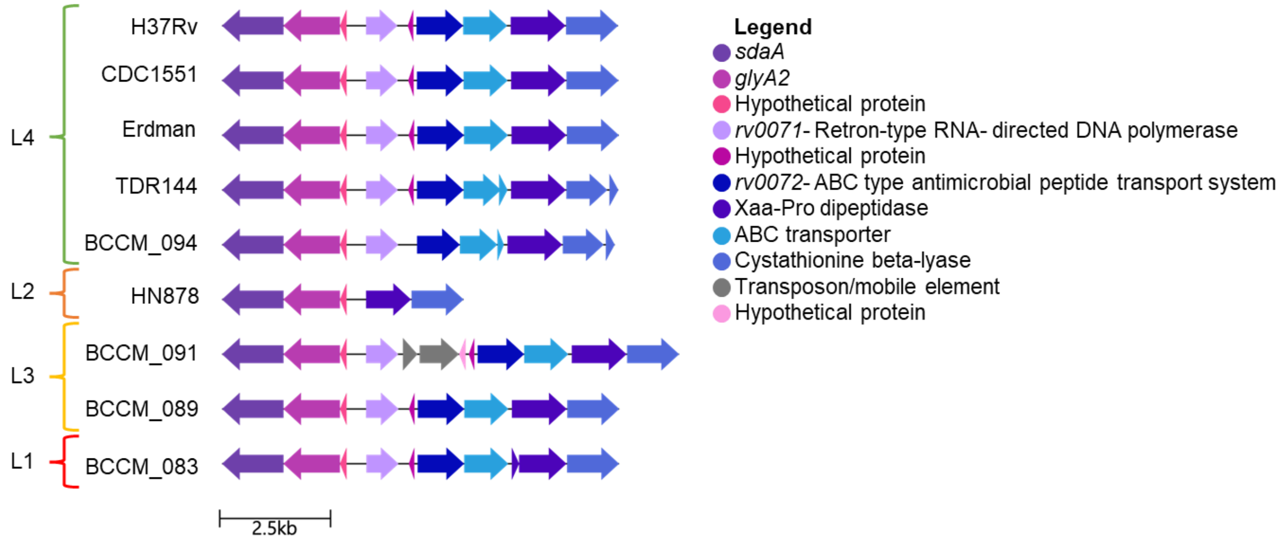

Fig. S3H

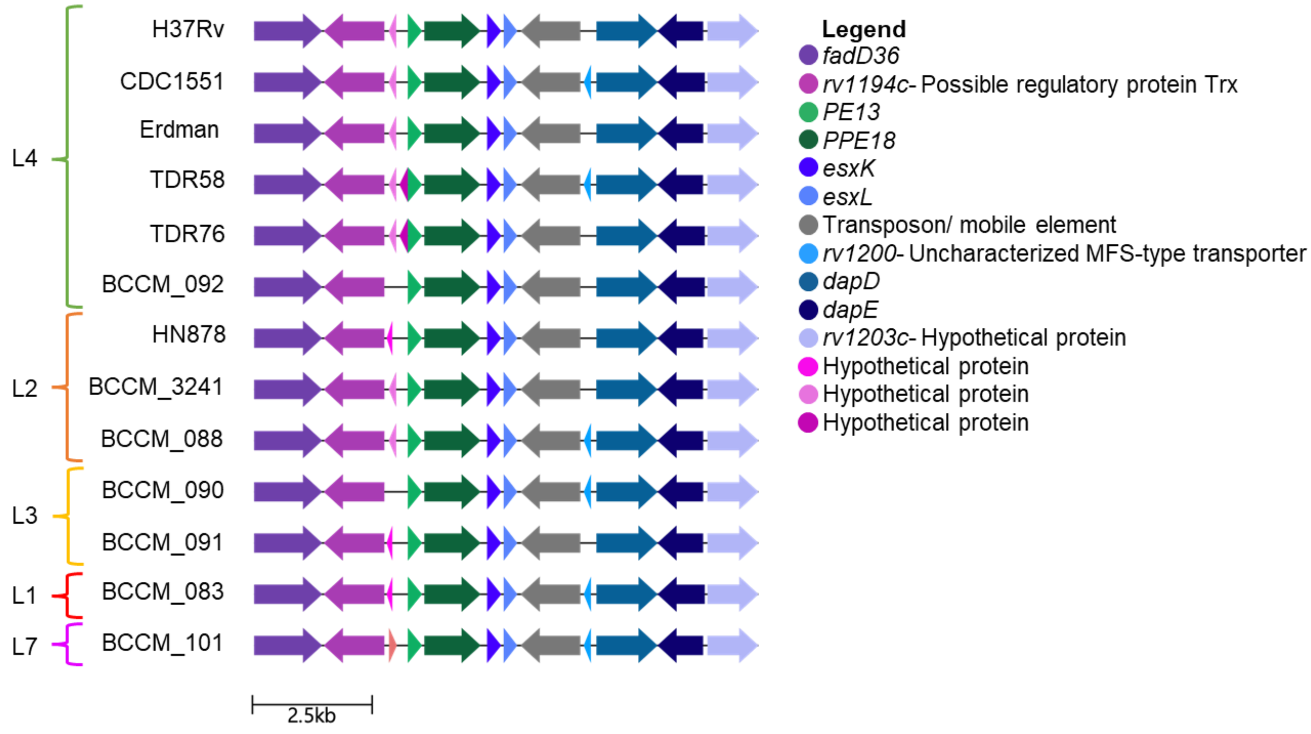

L4
H37Rv
CDC1551
Erdman
TDR58
TDR76
BCCM_092
L2
HN878
BCCM_3241
BCCM_088
L3
BCCM_090
BCCM_091
L1
BCCM_083
L7
BCCM_101
2.5kb
Legend
fadD36
rv1194c- Possible regulatory protein Trx
PE13
PPE18
esxK
esxL
Transposon/ mobile element
rv1200- Uncharacterized MFS-type transporter
dapD
dapE
rv1203c- Hypothetical protein
Hypothetical protein
Hypothetical protein
Hypothetical protein

### Fig. S3I

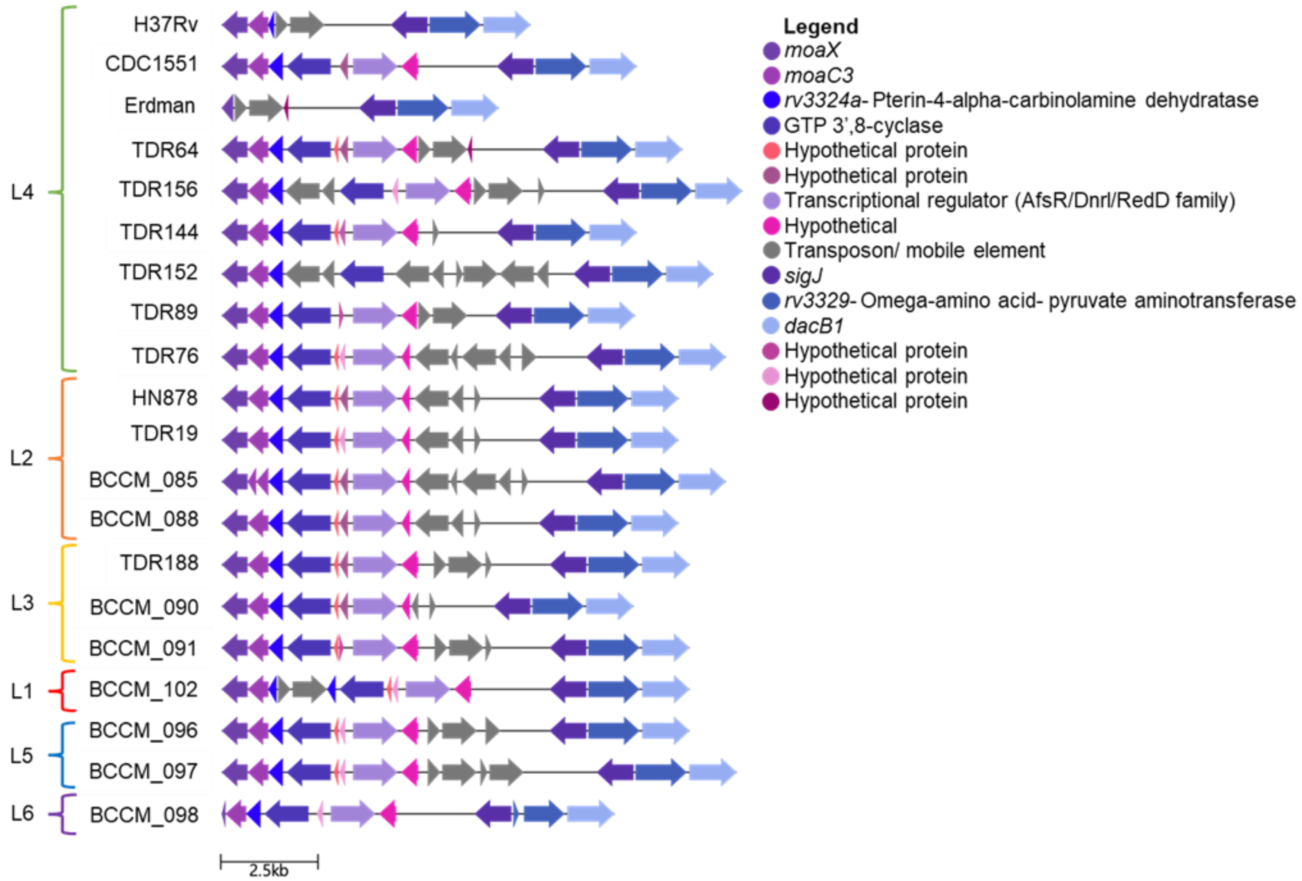

L4
L2
L3
L1
L5
L6
H37Rv
CDC1551
Erdman
TDR64
TDR156
TDR144
TDR152
TDR89
TDR76
HN878
TDR19
BCCM_085
BCCM_088
TDR188
BCCM_090
BCCM_091
BCCM_102
BCCM_096
BCCM_097
BCCM_098
2.5kb
Legend
moaX
moaC3
rv3324a- Pterin-4-alpha-carbinolamine dehydratase
GTP 3',8-cyclase
Hypothetical protein
Hypothetical protein
Transcriptional regulator (AfsR/Dnrl/RedD family)
Hypothetical
Transposon/ mobile element
sigJ
rv3329- Omega-amino acid- pyruvate aminotransferase
dacB1
Hypothetical protein
Hypothetical protein
Hypothetical protein

### Fig. S3J

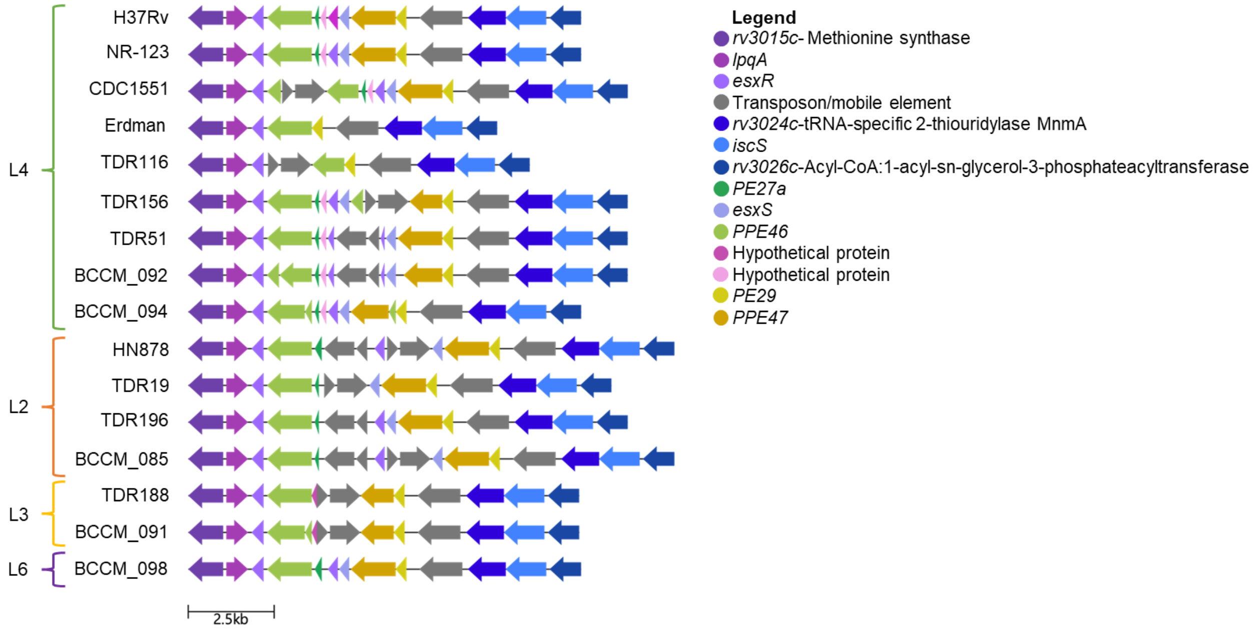


### Fig. S3K

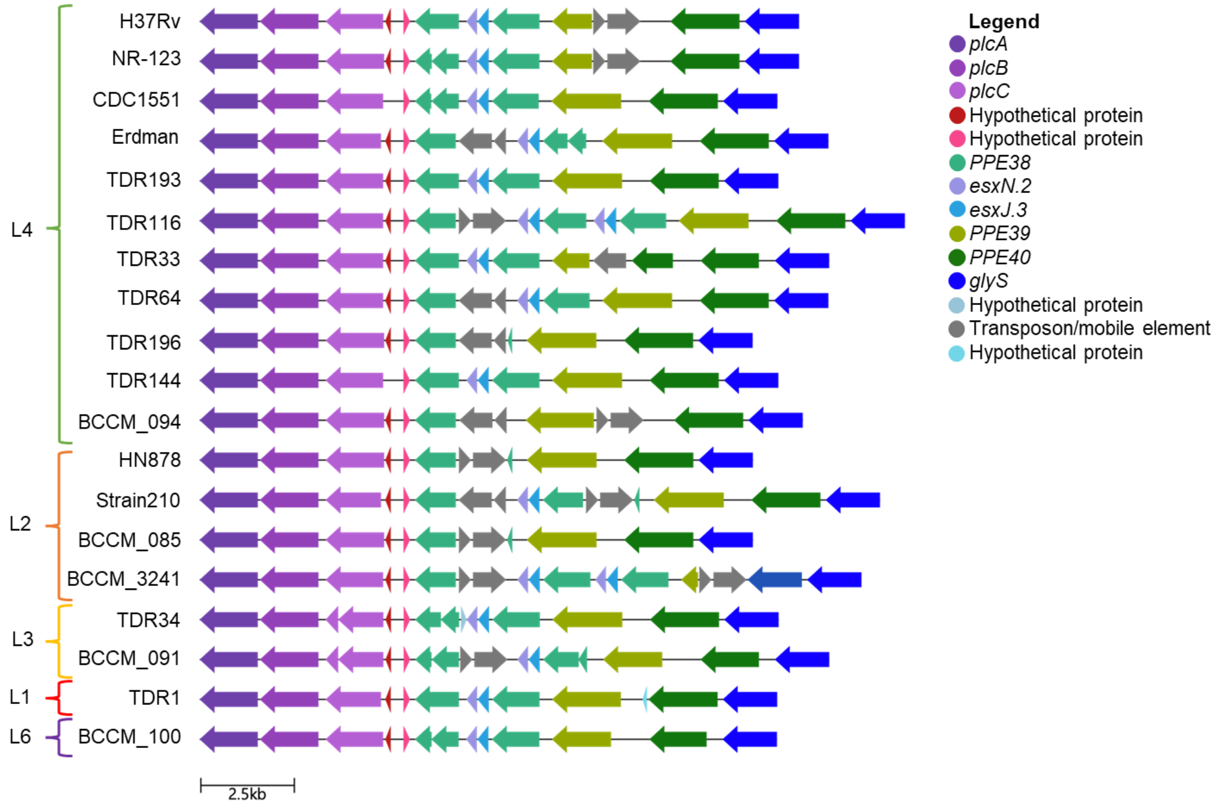

### Fig. S3L

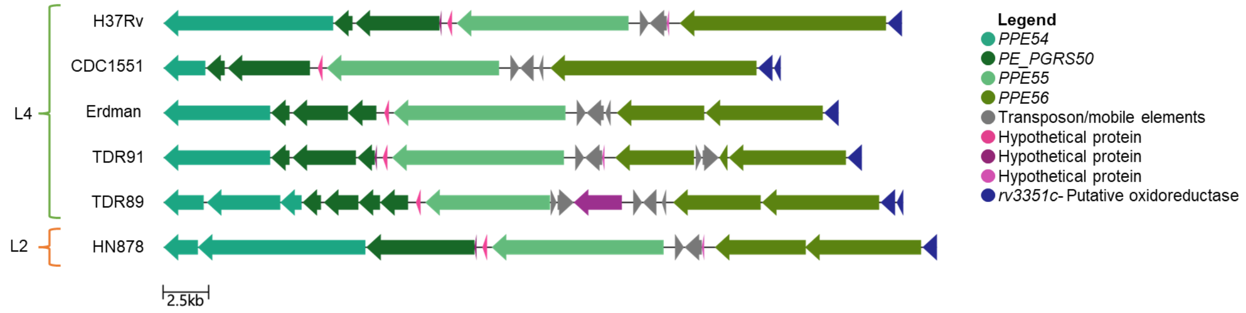


### Fig. S3M

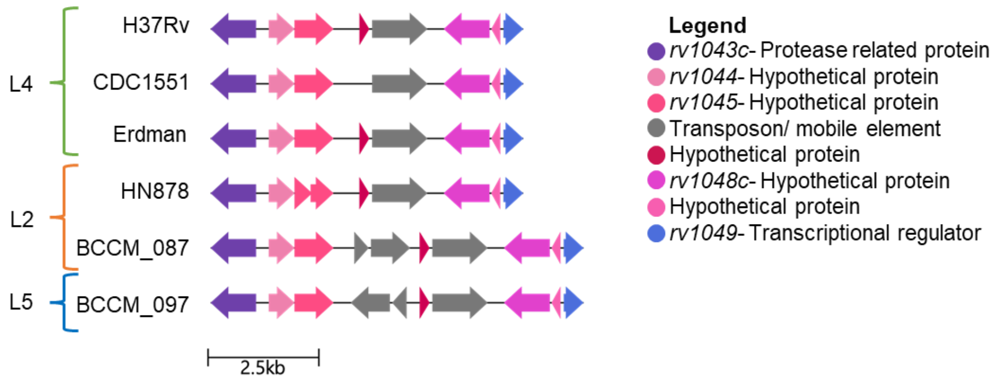


**Fig. S3. HVR regions across the M. tuberculosis Pangenome.** (A-M) Schematics of remaining HVR's not shown in main text as determined by PPanGGOLiN in representative strains and commonly used reference strains: H37Rv, CDC1551, Erdman, and HN878. Images were made using Clinker (57).

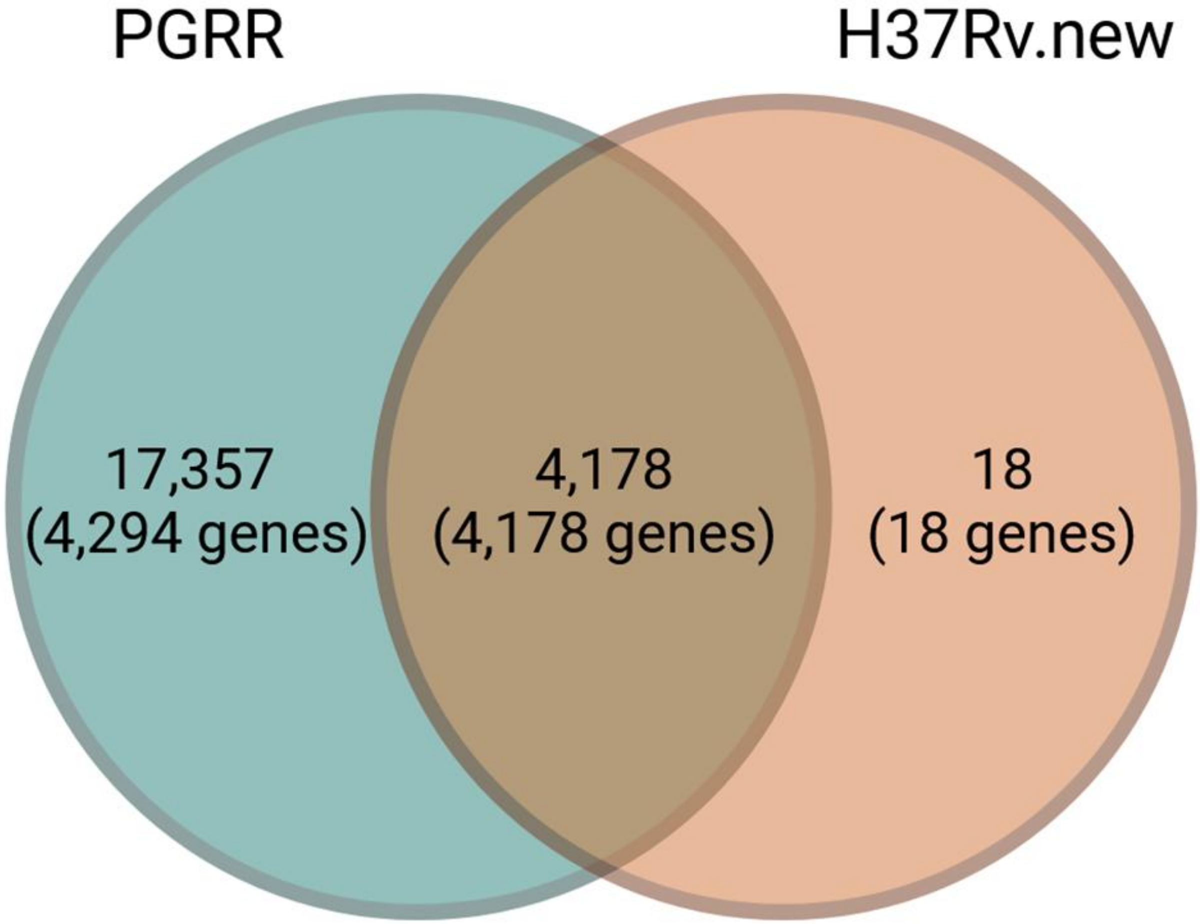


**Fig. S4.** Paralog alignment results of BCCM082 replicates to PGRR and H37Rv.new. Number of genes shown in figure are not exclusive.
