## Supplementary Methods for "Defining the *Mycobacterium tuberculosis* Pangenome and Suggestions for a New Composite Reference Sequence"

**Bacterial strains**. The 50 strains used in this study were predominantly sourced from 2 strain banks: The Special Programme for Research and Training in Tropical Diseases (TDR strain bank) which consists of 229 geographically diverse clinical *Mtb* strains [24] and a collection of 20 strains representing all seven lineages described previously [24, 25]. A complete list of strains and related information can be found in Supplementary Table 1.

**Pyrazinamidase-activity determination using a modified Wayne's method.** Agar slants were made by placing 7H9 1% w/v supplemented with 10% OADC and 400 ug/mL pyrazinamide (in DMSO) into agar tubes. Each *Mtb* strain to be tested was cultured to mid-log phase (OD595 = 0.6), spun down, and all but 150 uL of the supernatant discarded. The pellets were then resuspended and inoculated onto the surface of the agar slants and incubated at 37 C for 4 days. After incubation, 500 uL of freshly prepared 1% ferrous ammonium sulphate solution was added to each tube and incubated at 4 C for 4 hours. The color of the agar slants was then recorded. No color change indicated a negative test and a color change to orange-red indicated a positive test.

**Culture and DNA extraction.** *Mtb* strains from lineages 1-4 and 7 were cultured in Middlebrook 7H9 (BD Difco, Franklin Lakes, USA), 10% oleic acid-albumin-dextrose catalase (OADC) (BD Difco, Frankin Lakes, USA), 0.05% (vol/vol) Tween 80 (Sigma-Aldrich, St. Louis, USA) and 0.2% (vol/vol) glycerol (Sigma-Aldrich, St. Louis, USA). Strains from Lineages 5 and 6 share a conserved mutation in *pykA* [26] were cultured in media described above and supplemented with 40mM sodium pyruvate (Sigma-Aldrich, St. Louis, USA). Late exponential phase cultures were plated on 7H11 (Sigma-Aldrich,

St. Louis, USA) plates + 0.5% (vol/vol) glycerol (Lineages 5 and 6 strains were supplemented with or 40mM sodium pyruvate for) and extracted using Cetyltrimethylammonium bromide (CTAB) method [27] as described previously [13].

**Genomic DNA (gDNA) sample quality check**. After extraction the gDNA concentration and quality was evaluated using a Nanodrop One™ (ThermoFisher, Waltham, USA). Fragment size and DNA integrity numbers (DIN score) was evaluated using Genomic DNA Screen Tape on a 4200 TapeStation system (Agilent, Santa Clara, USA). The gDNA that met the following cutoffs were used for sequencing: A260/280 ratio: between 1.7-2.0; A260/230 ratio: between 1.5-2.5; DIN score $\geq$7; and DNA length >20 kb. Purified DNA was stored at 4°C prior to preparation of sequencing libraries.

**Whole genome sequencing (WGS**). *Nanopore Sequencing.* All gDNA was processed with a 1D sequencing kit (SQK-LSK109) (Oxford Nanopore Technologies (ONT, Oxford, United Kingdom) along with a native barcoding kit (EXP-NBD103 and EXP-NBD112) according to the native barcoding gDNA protocol. The gDNA was not sheared but used directly for DNA end repair and ligation. Both ligation steps in the protocol were extended from 10 min to 30 min. Six samples were multiplexed per run and the pooled adapter ligation step was performed in duplicate before sequencing. The final library was sequenced using a FLO-MIN106 flow cell on a MinION instrument.

**Illumina sequencing**. WGS Illumina reads were used to polish consensus sequences for all samples included in this study. All samples except for the BCCM samples were library prepped using an Illumina DNA PCR-free library prep kit (Illumina, San Diego, USA). Sequencing libraries were quality checked using the Qubit™ ssDNA kit (Thermo Fisher, Waltham, USA) and equal volumes were pooled for sequencing. The libraries were then sequenced in a paired-end 150 bp configuration on an Illumina NovaSeq 6000 platform. The 20 BCCM samples were paired-end sequenced on the Illumina HiSeq2500 with 101 cycles [25]. We obtained the raw sequencing results for the 20 previously sequenced BCCM strains from the European Nucleotide Archives (ENA) using the study accession number PRJEB27802 [25]. Supplementary Table 1 lists strain specific accession numbers.

**Assembly and annotation**. Strains were assembled using Bact-Builder (v.1.1, https://github.com/alemenze/bact-builder) [13]. The pipeline was designed for and optimized for the Rutgers High Performance Computing (HPC) cluster but has been restructured for use on any standard HPC or cloud-based resources. The local Rutgers (Amarel) cluster environment used for this analysis consisted of base nodes with 2x Intel Xeon Gold 6230 R (Cascade Lake) Processors (35.75 MB cache, 2.10 GHz): 2933 MHz DDR4 memory, 26-core processors (52 cores/node), 12×16 GB DIMMS (192 GB/node) per node, and GPU nodes included graphics cards with the NVIDIA Pascal architecture. All sequences used passes the Bact-Builder quality checks as described [13]. Complete genomes were annotated using RAST through the PATRIC interface [28-31].

**Lineage identification**. Illumina sequencing reads were analyzed using Mykrobe (Predictor version v0.10.0, Desktop app version v0.10.0) to confirm lineage predictions [32]. Lineage identification was confirmed by phylogenetic analysis generated by REALPHY (https://realphy.unibas.ch/realphy/) a reference sequence alignment based phylogeny builder [33].

**SNP and INDEL identification**. Final assemblies were assessed for SNPs, INDELs and regions of difference using DNAdiff (v1.3) from the MUMmer program (https://github.com/mummer4/mummer) [34] and implemented using CONCOCT (https://github.com/BinPro/CONCOCT) [35]. Significance of lineage SNP and INDEL differences relative to H37RV and intra-lineage SNP and INDELs were determined using pairwise t-tests adjusting using FDR multiple testing correction.

**Phylogenetic analysis**. A previously described whole genome multilocus sequencing typing (wgMLST) approach for *mycobacterium tuberculosis* complex strains was used for phylogenetic analysis [36] which has >97% concordance with previously defined lineage classifications [37]. This approach utilizes the 'Mycobacterium tuberculosis/bovis/africanum/canettii' schema available at http://www.cgmlst.org/mtbc for 2,891 core genes and 755 accessory genes. However, it excludes PE/PPE genes. The chewBBACA (v3.3.9) program, an allele calling tool that can be used with schemas from the cgMLST database, was used to determine the allelic profiles [38]. The AlleleCall module generated the allelic profiles from the set of 50 genomes, while the AlleleCallEvaluator module was used to generate a multiple-sequence alignment (MSA) and allelic distance matrix. A maximum likelihood tree was constructed from the MSA using IQ-TREE 2 (v2.2.2.7, http://www.iqtree.org/) [39], which inferred the best fit model Q.mammal+F+I. The tree was rooted at midpoint and the wgMLST tree plot was generated using R (v4.3.1) with ggtree (v3.8.2) [40] and ggplot2 (v3.5.1) [41].

**Pangenome analysis.** We identified core and accessory gene clusters across *Mtb* complex strains using the pangenomics workflow implemented in anvi'o (https://anvio.org) development branch of v8 [42, 43] with default parameters following the tutorial at https://merenlab.org/p/. The anvi'o pangenomics workflow enables the annotation of genes in genomes with functions using multiple databases. For our downstream interpretations of functions in core and accessory genes, we used the NCBI's Clusters of Orthologous Genes (COGs) database [44]. We interpreted the resulting pangenome using the anvi'o script anvi-compute-rarefaction-curves, which calculated and visualized rarefaction curves, and employed Heap's Law to compute the extent of closeness of the pangenome. A fully reproducible Mtb pangenome as a 'digital microbe' [45] is available via doi:10.6084/m9.figshare.28513640. To study what features were contributing to copy number variation in the core genome we generated an anvi'o pangenomics workflow based on a small cohort of 8 strains and generated a summary file of the core genome (Table S5). Using the python script "get_gene_cluster_names_with_unequal_num_paralogs_from_individual_genomes.py" we identified core gene clusters with copy number variation and studied a random subset of 25 gene clusters within our cohort. We used PPanGGOLiN v1.2.74 (https://github.com/labgem/PPanGGOLiN) [46] with default parameters to identify hypervariable regions. We assessed the differences in gene count and composition across lineages of *Mtb* and *M. canettii* using a one-sided *t*-test, and a Principal Component Analysis (PCA) plot illustrating Bray-Curtis dissimilarity distances on gene cluster presence-absence data reported for the pangenome by `anvi-summarize`, and used PERMANOVA to survey differences between lineages. R packages vegan v2.6.4 [47] and ggpubr v0.6.0 [48] created the PCA plot and gene count visualizations, respectively.

**Pangenome Gene Reference Resource**. We created a single resource that summarizes gene content and aggregates unique genes and their variants across our 50 *Mtb* strains **(https://github.com/wejlab/PGRR).** This Pangenome Gene Reference

Resource (PGRR) encompasses the complete panorama of genetic variation across genes defined by this study and can be used to map variability across genes of interest and align sequencing data for new strains. To create the PGRR, we extracted all genes across all pangenome strains and identified core and accessory genes whose sequences were perfectly conserved between strains. Each of these genes were represented once on the PGRR and labeled to point to all genomes that shared that exact gene sequence. Every unique paralog of these genes was also added to the PGRR and labeled to point to all genes that shared the exact sequence of that paralog. The completed PGRR contained 3,195 of these genes along with 21,870 of their paralogs. All duplicates and non-duplicated sequences were consolidated with their respective gene name and genome of origin into a single FASTA file (File S2).

**PGRR validation analysis**. Validation used DNA sequencing reads from *Mtb* strain BCCM082 (a Lineage 1 strain). To simplify this process, we first evaluated alignments over a known region of deletion called the *Mtb* specific deletion 1 (TbD1) region and then evaluated full genome alignment. Reads aligning specifically to the TbD1 region in BCCM082 were first aligned to the recently updated H37Rv reference (denoted as H37Rv.new in [13]), followed by alignment to the PGRR using Bowtie 2 with the "very-sensitive-local" parameter for all alignments. We then mapped all illumina sequencing reads from strain BCCM082 to the complete genome of strain BCCM082, H37Rv, and the PGRR using default "very-sensitive-local" Bowtie setting. Reads that did not align to H37Rv were re-aligned to the PGRR.
